# Lipoprotein(a) promotes thrombosis through platelet activation and promotion of a lysis-resistant thrombus architecture

**DOI:** 10.64898/2026.07.29.739869

**Authors:** Justin R. Clark, Frances S. Sutherland, Julia M. Assini, Arnaud Girard, Sébastien Thériault, Benoit J. Arsenault, Marlys L. Koschinsky, Michael B. Boffa

**Author notes:** Correspondence to: Michael B. Boffa, PhD, Room 4245C, Robarts Research Institute, The University of Western Ontario, 1151 Richmond St N, London, ON, Canada N6A 5B7.

## Abstract

Elevated levels of lipoprotein(a) (Lp(a)) are an independent risk factor for the development of atherothrombotic diseases. However, it is unknown if Lp(a) directly promotes thrombus formation, inhibits thrombus clearance, or merely accelerates the underlying atherosclerotic processes that culminate in plaque rupture. While numerous studies indicate that the apolipoprotein(a) (apo(a)) component of Lp(a) can inhibit plasminogen activation and fibrinolysis, recent evidence suggests that these effects may not be retained in Lp(a). An alternative mechanism through which Lp(a) may promote atherothrombotic events is by impacting platelet function. However, the effects of Lp(a) on platelet function and thrombosis have never been directly assessed in blood clots formed from flowing whole blood. Using a transgenic mouse model expressing high plasma concentrations of apo(a), we showed using a laser-induced mesenteric vessel injury model employing intravital microscopy that apo(a) increased platelet and fibrin volumes in the thrombi without affecting fibrinolysis. In a ferric chloride-induced mouse carotid artery thrombosis model, we found that apo(a) substantially reduced occlusion times and led to more stable thrombi; importantly, we also demonstrated that the effects of Lp(a) could be mitigated by low-dose aspirin therapy. We evaluated the prothrombotic potential of Lp(a) in human blood clots formed under arterial flow conditions using a Chandler loop apparatus. In these studies, we showed that the presence of Lp(a) during thrombogenesis inhibited lysis of the thrombi, without directly impacting fibrinolysis. Lp(a) promoted platelet accumulation in the Chandler thrombi and facilitated the development of fibrin networks that displayed features of fibrinolysis resistance. In an analysis of the UK Biobank, participants with Lp(a) ≥125 nmol/L had a higher risk for arterial thrombosis of non-atherosclerotic etiology but not a higher risk for venous thromboembolism. Collectively, these findings demonstrate that Lp(a) is inherently prothrombotic and likely promotes arterial thrombosis *in vivo* in part through promoting platelet activation. These findings explain how elevated Lp(a) is an important risk factor for arterial thrombosis either with or without an atherosclerotic etiology as well as observational primary prevention data suggesting that aspirin reduces atherothrombotic risk specifically in patients with elevated Lp(a).

## INTRODUCTION

Elevated plasma levels of lipoprotein(a) (Lp(a)) are well-established as an independent and causal risk factor for atherosclerotic cardiovascular diseases (ASCVD).^1^ However, there remain many unanswered questions regarding the mechanisms underlying Lp(a)-mediated pathophysiology.

Although Lp(a) is structurally similar to low-density lipoprotein (LDL), it is distinguished by the unique apolipoprotein(a) (apo(a)) that is covalently linked to apoB-100 in the LDL particle, and that bears striking sequence similarity to fibrinolytic proenzyme plasminogen.^2^

That Lp(a) is a potent cardiovascular risk factor likely reflects proatherosclerotic, proinflammatory, and prothrombotic properties of this complex lipoprotein.^3^ Its ability to preferentially bind to and transport highly inflammatory oxidized phospholipid species (oxPL) in the circulation to sites of arterial damage may underlie its role in the development and progression of atherosclerosis.^4–6^ Lp(a) impacts numerous vascular and immune cell types crucial to the atherosclerosis process, including endothelial cells^7,8^, smooth muscle cells^9^, monocytes^10–12^, and macrophages.^13–15^ Emerging data from imaging studies show that elevated Lp(a) is associated with vulnerable focal lesions marked by the presence of key high-risk plaque features such as increased thin fibrous cap atheroma, increased macrophage infiltration and larger lipid cores.^16–19^ Such plaques are prone to rupture leading to thrombosis and ischemic events. Data from our own transgenic mouse model for human Lp(a) likewise showed increased plaque area, increased necrotic core area, decreased collagen organization, increased oxPL deposition, and increased calcification of aortic atheromas.^20^

Lp(a) has also been suggested, more controversially, to possess prothrombotic properties.^21^ Initially, the homology of apo(a) to plasminogen was proposed to allow Lp(a) to compete for the physiological functions of plasmin(ogen) in fibrin clot lysis.^2^ However, while the apo(a) component of Lp(a) appears to display robust antifibrinolytic effects *in vitro*^22^, recent reports show that this is not the case for Lp(a). *Ex vivo* studies using plasma from patients in which dramatic reductions of plasma Lp(a) levels had been achieved using antisense oligonucleotide therapy showed no significant effect of Lp(a) lowering on plasma clot lysis times.^23^ We subsequently showed that the inability of Lp(a) to inhibit plasminogen activation and fibrinolysis results from masking of a key plasminogen-binding lysine residue in the protease-like domain of apo(a) when in the context of covalent attachment to apoB in the Lp(a) particle.^24^ However, in this study we also demonstrated that Lp(a) promotes an altered, lysis-resistant, fibrin clot structure.^24^ This finding is consistent with reports that found correlations between Lp(a) levels and reduced clot permeability^25^ and increased clot density.^26^ Moreover, we found that Lp(a) can accelerate clot formation and thrombin generation, the latter through effects on the contact pathway.^24^ It has also recently been demonstrated that monocytes from high-Lp(a) individuals expressed higher levels of tissue factor (TF) and that treatment of monocytes with purified Lp(a) stimulated increased TF expression.^11^

Together, these data pose a provocative question: does Lp(a) promote atherothrombotic events indirectly by promoting vulnerable plaques, or is Lp(a) itself inherently prothrombotic? Notably, it has been consistently reported that elevated Lp(a) does not increase risk for venous thromboembolism^27–30^, except at extremely high levels (i.e., above the 95th percentile).^29,31^ While confounding, these findings may reflect differences in clot composition in the venous and arterial systems. In general, arterial thrombi (‘white’ clots) contain fibrin and are platelet rich, while venous thrombi (‘red’ clots) are fibrin-rich and relatively platelet-poor.^32,33^ This has been interpreted to suggest a possible role for platelets in the atherothrombotic effects of Lp(a). Indeed, several *in vitro* studies show that Lp(a) enhances platelet aggregation in response to various agonists.^34–36^ Intriguingly, emerging clinical evidence suggests that individuals with high Lp(a) levels may specifically derive benefit (reduced major adverse cardiovascular events or cardiovascular mortality) from low-dose aspirin treatment in primary prevention settings.^37–39^ Furthermore, elevated Lp(a) levels have been shown to be associated with greater thrombus burden in younger individuals with ST-segment elevation myocardial infarctions^40^, and with left atrial thrombus formation in individuals with non-valvular atrial fibrillation.^41^

Nonetheless, direct evidence that Lp(a) is inherently prothrombotic is currently sparse. A caveat of many previous *in vitro* or *ex vivo* studies assessing the role of apo(a) and Lp(a) in thrombosis is the use of plasma rather than whole blood and using static *in vitro* assays as opposed to clots formed under flow. Moreover, studies of a prothrombotic role for Lp(a) in animal models are completely lacking. In the present report, we address these knowledge gaps using two different thrombosis models in transgenic mice expressing human apo(a), and studies of the structure and propensity to lyse of thrombi formed from whole blood under flow in a Chandler loop. We have also performed a novel analysis of UK Biobank data to support the idea that elevated Lp(a) promotes arterial thrombosis in non-atherosclerotic contexts.

## MATERIALS AND METHODS

### Transgenic LPA mouse model for *in vivo* thrombosis studies

Details for the generation and genotyping of mice expressing the human *LPA* construct have been previously described.^42^ Briefly, the *LPA* transgene was constructed in the pLIV.7 vector and encodes a 13K form of apo(a). The transgene was maintained in the hemizygous (*LPA*^+/0^) state and crossed with non-transgenic mice of the same background strain to yield transgenic apo(a)-expressing (*Tg*(*LPA*^+/0^); *LPA*) and wild-type control (*Tg*(*LPA*^0/0^); *WT*) progeny. Expression of the *LPA* transgene was verified by western blot analysis of mouse plasma. Plasma apo(a), cholesterol, and triglyceride measurements in mouse plasma were determined using Randox reagent kits (apo(a): LP3403; total cholesterol: CH200; and triglyceride TR210) and a Randox Daytona clinical autoanalyzer according to the manufacturer’s instructions. Approval for breeding and experimental animal procedures was obtained from the Animal Care Committee at Western University (AUPs 2020-104 and 2020-107, respectively).

Five- to six-week-old male and female mice were fed *ad libitum* either a standard diet (TD8604, Envigo) or a high-fat, high-cholesterol diet (HFHC: 42% calories from fat, 0.2% cholesterol; TD88137, Envigo) for 5 to 6 weeks. The mice were also injected weekly with 5 mg/kg (intraperitoneally (i.p.)) of a GalNAc-conjugated antisense oligonucleotide (ASO) targeting the mouse *Ldlr* gene encoding the LDL-receptor (provided by IONIS Pharmaceuticals).^20,43^ The HFHC diet and *Ldlr* knockdown results in substantial hyperlipidemia (**Supplementary Fig. S1**), in keeping with our previous studies.^20^ Mice were housed at 23°C on a 12-h light/dark cycle, and weight gain was monitored weekly. Following 5 to 6 weeks of feeding and weekly *Ldlr* ASO injections, the mice underwent thrombosis experimental protocols; 4 – 6 mice of each genotype and sex were used in all cases. In some experiments, a subset of mice also had their daily *ad libitum* water supplemented with acetylsalicylic acid (aspirin; Millipore Sigma) during the last two weeks prior to experimental procedures to achieve an intake of 25 mg/kg/day. It has previously been shown that this dose of aspirin in mice has similar pharmacodynamic effects to low-dose aspirin treatment in humans.^44^ Water was replaced every other day to avoid precipitation of the aspirin, and intake was monitored daily.

Following completion of the experimental protocol, cardiac puncture was conducted to collect blood into syringes containing 7% Na_2_-EDTA; blood was then centrifuged for 12 min at 12,000 × *g*, 4°C to obtain plasma.

### Intravital imaging of laser-induced thrombosis

Intravital microscopy and laser-induced endothelial injury procedures were adapted from a previously described protocol optimization.^45^ Mice were anesthetized in an induction chamber using 3.5% (v/v) isoflurane mixed with 100% medical grade oxygen at 0.5 L/min (SomnoSuite Low-Flow Anesthesia System). The anesthetized mouse was then placed onto a heating pad, and a nose cone was fitted to ensure maintenance of isoflurane anesthesia (2% (v/v) + 100% medical-grade oxygen at a 0.5-1 L/min flow rate). After an appropriate depth of anesthesia was reached, a midline incision was created, and the mesentery was exposed. The exteriorized mesentery was moistened with warm sterile saline, and then fluorescently labelled anti-platelet GPIb*β* (Emfret DyLight-649; 0.1 µg/g bodyweight) and anti-fibrin antibodies (Millipore Sigma; 0.2 µg/g bodyweight; labelled using Alexa-Fluor-546 labelling kit) were intravenously injected into the inferior vena cava. While maintaining anesthesia via nose cone ventilation, the mouse was placed on its side onto the microscope stage. A heating pad was used to maintain thermal regulation. The intestines were then gently teased apart and flattened onto the microscope stage using cotton tips wetted with sterile saline. For consistency, mesenteric vessels in the ileum region were used for data collection.

Mesenteric veins (MV) with an outer diameter range of 300-700 µm were selected as targets for laser injury.^45^ After locating the endothelium in a target MV using auto-fluorescent elastin (488-nm) as a guide, a Nikon A1R+ confocal microscope equipped with a 50 mW 405 nm laser was used for ablation of a 75 × 75 µm region of interest (ROI) within the endothelial layer. For all image acquisitions, a 20× dipping water-immersion lens objective was used. Our microscope was also equipped with a motorized Piezo Z-stage for fast z-stack acquisitions. Images collected were 512 × 512 pixels, and emitted fluorescence signals in the 488-nm (elastin), 561-nm (fibrin) and 647-nm (platelets) channels were detected simultaneously. Z-stack images with a step size of 1 µm and a z-range of 87 µm (−17 µm to +70 µm) were acquired, which enabled capture of the entire depth of the resultant thrombus. Prior to endothelial injury, a “pre-injury” z-stack, using the same settings mentioned, was acquired to visualize vessel and tissue architecture before the injury commenced. Following injury, z-stacks were collected every minute for 17 min. At least 1 injury was made per mouse. In situations where multiple injuries were made on the same mouse, only neighboring MVs or regions upstream of a previous injury within the same MV were used.

All files were subjected to identical image processing, threshold settings and analysis steps using the NIS Elements Advanced Research Software (Nikon). Images were cropped, denoised, thresholded, and processed using the smoothing and cleaning features of the software to separate the thrombus mass from flowing platelets in the vessel lumen. Next, platelet and fibrin volumes were quantified by the imaging software at each individual timepoint (1-17 min). In situations where 2 injuries were introduced on a single mouse, platelet and fibrin volumes were averaged at each timepoint for that animal. Platelet and fibrin volume Area Under the Curve (AUC) measurements were compiled.

### Ultrasound flow-probe analysis of FeCl_3_-induced carotid artery injuries

The ferric chloride (FeCl_3_)-induced carotid artery injury model was used to assess rates of occlusive clot formation.^46–48^ Briefly, on the day of the experiment, anesthetized mice were placed onto a heating pad, and a nose cone was fitted to maintain isoflurane delivery (2-3% (v/v) + 100% medical-grade oxygen at a 0.5-1 L/min flow rate). When an appropriate depth of anesthesia was achieved, the right common carotid artery (RCA) was exposed by blunt dissection. Care was taken to separate the vagus nerve from the RCA. A miniature Doppler flow probe (0.5PSB, Transonic Systems Inc.) was then fitted to the RCA to monitor blood flow, and baseline flow measurements were recorded.

Thrombus formation was induced by placing a piece of filter paper (1 mm × 2 mm) soaked in 30% FeCl_3_ (anhydrous; equals 1.85 M) directly in contact with the adventitial surface of the vessel. The filter paper was placed onto the RCA in a region upstream of the flow probe (towards the direction of the heart) for 2.5 min (male mice) and 1.5 min (female mice). Following injury, the vessel was washed with sterile saline. Blood flow was continuously monitored using the flow probe from the time of baseline flow measurement, through the injury period, and up until a maximum of 15 min following the FeCl_3_-induced injury.

The endpoints for this experiment were (1) when blood flow had decreased to 25% of baseline flow (these clots were considered to be 100% occlusive) or (2) if occlusion was not seen within 15 min after injury. For the latter, 15 min was used as the value for statistical analyses. The number of embolic events observed per 15 min Doppler recording was also recorded. An embolic event was identified as a sudden spike in blood flow during a period of general flow decline (**Supplementary Fig. S2**).

### Purification of recombinant apo(a) variants and Lp(a)

Recombinant apo(a) variants including (1) a wild-type 17 Kringle (17K) apo(a) construct consisting of 8 KIV2 repeats with all other kringle subunits of apo(a) intact, (2) a 17K apo(a) variant lacking the strong lysine binding site (sLBS) in KIV10 (17KΔLBS10), and (3) a 17K apo(a) variant lacking the protease-like domain (17KΔP) have been previously described.^49^ All r-apo(a) variants were purified from conditioned medium of stably expressing HEK-293 cell lines using lysine-Sepharose affinity chromatography^49^; isolated proteins were assessed for purity by SDS-PAGE and Imperial staining. Lp(a) was purified, with informed consent, from the plasma of a single human donor by sequential flotation followed by ion exchange chromatography as previously described.^15^ The donor expresses a single 16K apo(a) isoform. Human experimentation was approved by the University of Western Ontario’s Research Ethics Board (REB# 118593).

### *In vitro* formation of thrombus analogues under conditions of arterial flow using a Chandler loop device

Citrated whole blood collected (with informed consent; REB# 118593) from a human donor with low Lp(a) levels was combined with 0.05 mg/mL (final concentration) of Alexa FluorTM 488-conjugated, plasma-derived human fibrinogen (Invitrogen). Next, the citrated blood was supplemented with 250 nmol/L of either plasma purified Lp(a), 17K wild-type apo(a), 17KΔLBS10, or 17KΔP, with HEPES buffered saline (HBS) as the vehicle control, and incubated at 37°C for 30 min. Directly prior to the start of the procedure, each sample was recalcified by addition of CaCl_2_ (10 mmol/L final), gently mixed, and then pipetted into a piece of polyvinylchloride (PVC) tubing (inner diameter = 3 mm, external diameter = 4.2 mm, length = 33 cm) in a final volume of 1 mL. Each end of the PVC tubing was end-joined using a short sleeve of larger tubing (internal diameter = 4 mm, external diameter = 6.8 mm) to create a Chandler loop with a radius of curvature of 5.25 cm. The PVC tubing containing blood mixtures were then rotated for 90 min at ambient temperature, under a calculated wall shear rate of approximately 550 s^-1^. This wall shear rate was selected since whole blood clots formed under these conditions have been shown to resemble arterial thrombi.^50–52^

Following the 90-min rotation, resultant clots (∼1 cm in length) were collected and washed in 0.9% (w/v) saline; the washed clots were weighed using an analytical balance, then added to microcentrifuge tubes containing autologous low-Lp(a) donor plasma supplemented with tissue-type plasminogen activator (tPA; 50 nmol/L final) to induce clot lysis. Plasma tubes containing thrombi and tPA were then incubated at 37°C and samples (10 µL) of the plasma mixture were removed at 30 min intervals up to 180 min. Each sample was added to a tube containing 240 µL PBS, mixed, and then fluorescence (representing fibrinolysis) was measured using a fluorescence plate reader at excitation: 493 nm and emission: 520 nm. To account for differences in fluorescence release due to variations in clot sizes, fluorescence readings for each variant groups were normalized using a weight correction factor, derived from the average recorded weights of clots (n=4) formed in the presence of that variant (**Supplementary Fig. S3**).

In some cases, clots formed in the presence of only HBS vehicle were placed into autologous plasma containing tPA, in the absence or presence of 250 nmol/L of either Lp(a), 17K, 17KΔLBS10, or 17KΔP, for analysis of tPA-mediated fibrinolysis. In other cases, a subset of thrombi formed in the presence of vehicle control or 250 nmol/L of either Lp(a), 17K, 17KΔLBS10, or 17KΔP were immediately fixed in 4% paraformaldehyde for 24 h, paraffin embedded, and longitudinally sectioned for histology and immunostaining analysis, as detailed below. A final subset of thrombi, formed in the presence of vehicle control or 250 nmol/L of either Lp(a) or 17K, was subjected to scanning electron microscopy imaging and analysis as detailed below.

### Histology and immunofluorescence staining of Chandler loop thrombi

Following fixation and embedding as described above, Chandler loop thrombi were longitudinally sectioned in 5-µm increments. Sections were retrieved from the longest longitudinal region of the clot, ensuring that the middle region of the clots was sufficient for immunofluorescence (IF) analysis. Ten to twelve sections were retrieved from each clot, and each section was deparaffinized. The first section of each clot was subjected to hematoxylin and eosin (H&E) staining; sections were scanned using a Leica Aperio AT2 bright-field digital slide scanner with a 40× objective (Aperio Technologies) For the remaining sections, antigen retrieval with Citrate Buffer pH 6.0 (Millipore Sigma; #C9999) and immunostaining was then performed. Following antigen retrieval, tissue sections were blocked with 10% normal serum and 1% bovine serum albumin (BSA; BioShop) in Tris-buffered Saline (TBS; 20 mmol/L Tris pH 7.4, 150 mmol/L NaCl) for 2 h. Sections were then co-incubated overnight at 4°C with an anti-apo(a) primary antibody (either rabbit monoclonal, Abcam #ab208184; 1:200, or an in-house mouse monoclonal antibody) as well as either an anti-fibrin primary antibody (mouse monoclonal, Millipore MABS2155; 1:250), anti-platelet (CD42b; GPIb) primary (mouse monoclonal, Beckman #IM0409; 2 µg/ml), or anti-Plasminogen Activator Inhibitor-1 (PAI-1) primary (rabbit polyclonal, Abcam #ab66705; 1:100). Slides were next washed twice in TBS containing 0.025% Triton X-100 and incubated with an Alexa-Fluor 647-conjugated donkey anti-rabbit IgG secondary antibody (Invitrogen #A31573; 1:250) or an Alexa-Fluor 488-conjugated donkey anti-mouse IgG secondary antibody (Invitrogen #A21202 1:250) for detection. In some cases, slides were also counterstained with DAPI (Thermo Fisher Scientific #D1306; 1:3000) to visualize nuclei. Stained sections were mounted to glass coverslips using fluorescent mounting medium (Dako).

### Confocal immunofluorescence microscopy

Stained thrombus sections were imaged using a Nikon AIR+ Confocal Laser Scanning System. For all slides, denoised tiled images of the entire thrombus were collected using a 20x water-immersion objective (512 × 512 pixels, 1.5 digital zoom) with simultaneous detection in the 405, 488, and 647 channels. The Nikon Elements ‘Focus Surface’ program was used to predict the appropriate focal height of all areas imaged on the slide during tiling. Denoised regions of interest were also collected using either a 40x oil-immersion objective (1024 x 1024 pixels, 13 z-slices, 0.925-µm step size, 3× digital zoom) or a 60x oil-immersion objective (1024 x 1024 pixels, 21 z-slices, 0.2-µm step size, 1× digital zoom) with simultaneous detection in the 405, 488, and 647 channels. For all representative images of the same target, identical laser power and gain settings were used.

### Immunofluorescence image analysis

QuPath software was used to quantify total clot areas and areas of IF-positive staining of each 20× tiled image. The area of positive staining was then expressed relative to the total clot area. Percent area of fibrin staining was determined using 6 z-stack images per thrombus, acquired throughout the fibrin-rich tail region of the clots with the 40× oil-immersion objective to better distinguish the network features of the fibrin.^50^ To determine the percent area of fibrin positive staining, ImageJ software (v2.14.0) was used to reduce noise using a Gaussian blur, and then images were binarized using the automated threshold Otsu method. Next, original images and binarized images were added using the AND function of ImageJ. Images were thresholded and the ‘Analyze Particle’ function was used to determine the percent surface area coverage of fibers per each slice in the z-stack. Fibrin pore diameters were determined using a previously described method.^53^ Pore diameter measurements were obtained from the same confocal images used to determine percent areas of fibrin staining. Briefly, maximum intensity projections were created using the z-project feature on ImageJ. Images were then binarized using the automated threshold Otsu method, and a 600-µm^2^ grid was overlaid to split the image into 6 sextants. Next, the line tool was used to manually measure the pore sizes. Three different diameter measurements were acquired for each pore and then averaged. In total, 3 randomly selected pores were measured within each sextant (18 total pores per each individual image). The pore diameters were averaged within each square, within all squares counted in a singular image, and then across all images for each replicate.

### Ultrastructural SEM analysis of Clots formed in the Chandler Loop

Chandler loop clots used in SEM experiments were formed in a similar manner as described above, with slight modifications. Specifically, low-Lp(a) donor blood was supplemented with either vehicle, or 250 nmol/L of either Lp(a) or 17K, added to a closed loop of PVC tubing, and then rotated at 550 s^-1^ for 40 min at 37°C to maximally preserve cell morphologies. Formed clots were carefully washed 3× for 5 min in deionized water, fixed in 2% (v/v) glutaraldehyde in 50 mM cacodylate buffer pH 7.4 containing 150 mM NaCl for 24 h, and then briefly rinsed. Following fixation, thrombi were cut longitudinally and then dehydrated in a series of immersions in increasing v/v concentrations of ethanol (30%, 50%, 70%, 80%, 90%, and 100%). Samples were then mounted onto aluminum stubs, with the interior of the clot facing upwards for imaging. Samples then underwent further dehydration in a 37°C oven for a minimum of 2 h before coating with osmium. High-definition micrographs were obtained at 2000× magnification using a Leo 1530 Scanning Electron Microscope at 3-kV accelerating voltage (Western Nanofabrication Facility). Three independent images were acquired from 5 random interior regions of each clot. Examination and quantitative analyses of the predominant structures within the samples were conducted as previously described.^33^ Briefly, a 1.5 µm × 1.5 µm square grid was overlaid onto each micrograph and then each structural component was quantitated as the number of grid squares occupied by that component for each image. The grid size was selected such that only 1 predefined structural element was present within each square. If a particular square was occupied by two structures in the same ratio, then each component was quantitated as a 0.5 structure. The sum of all squares occupied by each component was then divided by the total number of squares counted per image to provide a percentage area occupied by each structural component in that image. The identifiable structures within the Chandler loop clots were similar to what has been described in both murine thrombi, human arterial and venous thrombi, and pulmonary emboli.^33,54,55^ The following predefined structural elements^33^ were identified during quantitative analysis: (i) individual fibrin fibers resembling plasma clots; (ii) fibrin bundles, which are made up of tightly packed fibrin fibers and have been associated with clot contraction; (iii) fibrin sponge, which is made up of tightly packed fibrin fibers and has been shown to be inversely associated with thrombus resolution^54^; (iv) platelets, which contain both individual highly activated platelets with visible membrane extensions and platelet aggregates; (v) cellular microvesicles, which were identified as round structures with a diameter of less than 1 µm; (vi) biconcave red blood cells (RBCs); (vii) intermediate RBCs; (viii) polyhedral RBCs (polyhedrocytes); and (ix) empty space / other.

### Statistical analysis

In all cases, analyses were performed by observers blinded to the genotypes and sexes of mice or the experimental group of the thrombi. Statistical analyses were performed using GraphPad Prism (Version 10.2.3) software. Pairwise comparisons were conducted using a two-tailed Student’s *t*-test assuming unequal variances. Comparisons between multiple groups were conducted using one-way or two-way ANOVA, as appropriate, with Tukey’s post hoc test. Kaplan-Meier plots were compared using the log-rank test. Statistical significance was defined as p values less than 0.05.

### UK Biobank analysis of the relationship between Lp(a) levels and arterial versus venous thrombosis events

The UK Biobank is a population-based study in the United Kingdom that included more than 500 000 participants aged between 40 and 69 years recruited between 2006 and 2010 using the National Health Services’ central registers (NHS). Data collection included a detailed self-report lifestyle questionnaire that was verified by trained medical personnel, physical measures, and the collection of blood, saliva, and urine samples. These tests were conducted in 22 assessment centers during the recruitment period. UK Biobank received approval from the British National Health Service, North West - Haydock Research Ethics Committee (16/NW/0274). All participants provided signed and informed consent for their participation in this study. Lp(a) levels were measured using an immunoturbidometric assay on a Randox AU5800. The current analyses were conducted under UK Biobank data application number 25205.

The primary outcome studied was the incidence of arterial thrombosis or embolism. This outcome was defined using the ICD-10 code I74, which denotes the incidence of arterial embolism or thrombosis. The incidence of venous thromboembolism, a composite of deep vein thrombosis, pulmonary embolism, or venous thrombophlebitis was defined using ICD-10 I80.2, I82.2, I26.0, and I26.9, as well as OPCS-4 codes L79.1 and L90.2. Participants with diagnostic codes of either of these outcomes before their baseline assessment visit were excluded. Patients with an incidence of both arterial and deep vein thrombosis (n=216) were also excluded. Maximum follow-up time was set to 15 years.

Multivariable Cox proportional hazard ratios adjusted for age, sex, ethnicity, LDL-C levels, smoking status, body mass index (BMI), statin use, anti-hypertensive medication use, and insulin use were used to evaluate the effect of Lp(a) on the incidence risk of arterial embolism or thrombosis and venous thromboembolism. Hazard ratios (HR) and corresponding 95% confidence intervals (95% CI) were obtained. The Cox proportional hazard model assumptions were verified through visual inspection of the Schoenfeld residuals. Resulting fits were evaluated based on the concordance of the predicted models with the observed data. All statistical analyses were performed using the R computational software version 4.1.3 in conjunction with the packages “tidyverse”, “data.table”, “lubridate”, “survival”, “survminer”, “gt”, “gtsummary”, and “writexl”. Figures were created using the “ggplot2” package.

## RESULTS

### Apo(a) increases platelet and fibrin volumes following laser-induced thrombosis

Transgenic apo(a)-positive (Tg(*LPA*^+/0^); *LPA*) and wild-type apo(a)-negative (WT(*LPA*^0/0^); *WT*) mice were fed a HFHC diet with weekly *Ldlr* ASO injection for 5 to 6 weeks, then underwent laser-scanning endothelial injury to evaluate the impact of apo(a) on platelet and fibrin accumulation during thrombogenesis *in vivo*. Generally, we observed that laser injuries resulted in a prominent core-in-shell architecture where fibrin localized to the interior of the clot adjacent to the site of endothelial damage (the core) and was surrounded by a thick outer shell of platelets (**Fig. 1A,B,E,F; Supplementary Fig. 4A,B,E,F**). This observation is consistent with previous findings using a similar laser-inducing injury model in mice as well as with a growing body of evidence reporting that a core/shell architecture is also a common feature of acute ischemic stroke thrombi.^56–58^

**Fig. 1.**
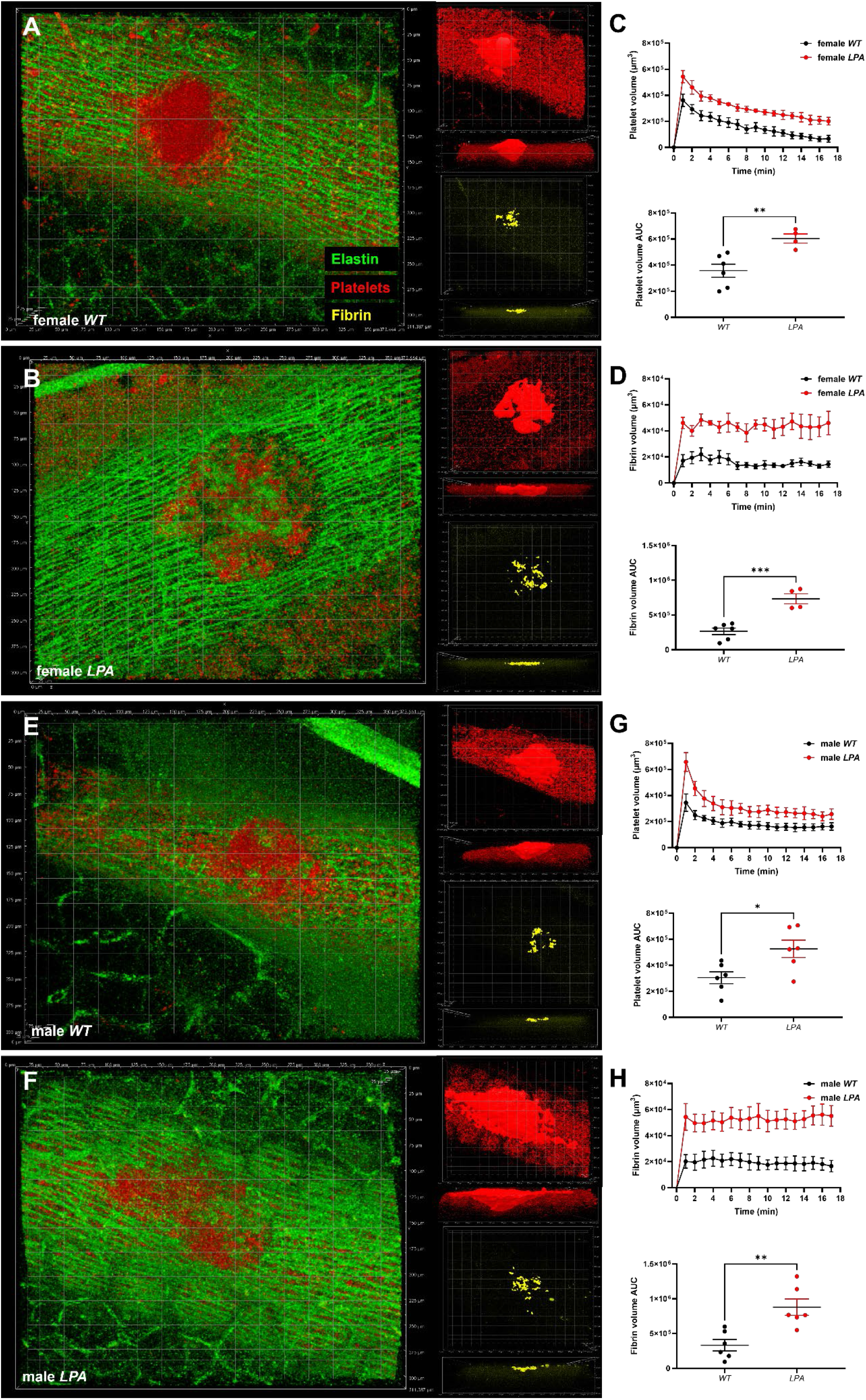
Apo(a) increases platelet and fibrin volumes in HFHC-fed mice in a laser-induced mesenteric vessel thrombosis model assessed by intravital imaging. Thrombus formation in mesenteric veins was followed for a period of 17 minutes to determine platelet and fibrin volumes within developing thrombi. Representative XY and XZ 3D surface reconstructions of thrombi formed at t=1 min in (**A**) wild-type (*WT*); n=5) and (**B**) transgenic apo(a)-expressing (*LPA*; n=6) female mice fed a HFHC diet. Overlay images contain colour composite green (emission wavelength: 488 nm; auto-fluorescent elastin), yellow (emission wavelength: 546 nm; fibrin), and red (emission wavelength: 647 nm; platelets). (**C**) Quantitative data displaying platelet volumes (upper panel) and scatterplots of distributions of cumulative platelet volume AUC (lower panel) in HFHC-fed female mice. (**D**) Quantitative data displaying fibrin volumes (upper panel) and scatterplots of distributions of cumulative fibrin volume AUC (lower panel) in HFHC-fed female mice. Representative XY and XZ 3D surface reconstructions of thrombi formed at t=1 min are also shown for (**E**) male *WT* (n=6) and (**F**) male *LPA* (n=5) mice fed a high fat/high cholesterol diet. (**G**) Quantitative data displaying platelet volumes (upper panel) and cumulative platelet volume AUC (lower panel) in HFHC-fed male mice. (**H**) Quantitative data displaying fibrin volumes (upper panel) and cumulative fibrin volume AUC (lower panel) in HFHC male mice. Quantitative data are shown as means ± SEM. Overlaid grid squares have dimensions of 25 × 25 µm^2^. \**p* < 0.05, \*\**p* < 0.01, \*\*\**p* < 0.001 by Student’s t-test.

In the HFHC-fed mice, there were substantial increases in both platelet and fibrin volumes in *LPA* mice versus *WT* mice throughout the 17-minute data collection period, in both males and females (**Fig. 1**). Accordingly, both mean platelet and fibrin areas under the curve (AUC), were significantly increased in the *LPA* mice compared to the *WT* mice (**Fig. 1G,H**).

In the standard diet-fed female mice, although a slight trend towards increased platelet and fibrin volumes was observed (**Supplementary Fig. 4C,D**), there were no significant differences in the respective AUC. In the standard diet-fed male mice, on the other hand, both platelet and fibrin volumes were notably increased in the *LPA* mice compared to *WT* controls and both mean platelet and fibrin areas under the curve (AUC) were significantly increased in the *LPA* mice compared to the *WT* mice (**Supplementary Fig. 4G,H**). The difference between male and female standard diet-fed mice and the greater effect size in HFHC diet-fed mice can likely be explained by the achieved plasma apo(a) levels in the respective mice: the apo(a) levels in HFHC diet-fed mice were 199.9 nmol/L and 83.8 nmol/L in males and females, respectively, and the corresponding values in standard diet-fed mice were 74.3 nmol/L and 34.7 nmol/L (**Supplementary Fig. S1**).

### Apo(a) enhances rates of occlusive clot formation following FeCl_3_-induced arterial thrombosis through a mechanism that is sensitive to aspirin administration

To further examine the role of Lp(a) on thrombogenesis *in vivo*, as well as evaluate the relationship between elevated apo(a), thrombosis, and low-dose aspirin therapy, 5- to 6-week-old transgenic *LPA* and control *WT* mice were fed HFHC diets with weekly *Ldlr* ASO injection for 5 to 6 weeks and then subjected to FeCl_3_-induced right carotid artery (RCA) injury. Downstream blood flow was monitored using a miniature Doppler flow probe to determine rates of embolic events and time to occlusive clot formation. A subset of mice also had their daily *ad libitum* water supplemented with low-dose aspirin (25 mg/kg/day) for 2 weeks prior to experimental procedures. Representative raw Doppler read-outs displaying embolic events and cessation of blood flow following injury are shown in **Supplementary Fig. S2**. Following FeCl_3_-induced injury, we observed a significant reduction in the mean occlusion time in the female *LPA* mice compared to the female *WT* mice (4.63 min versus 8.98 min; p < 0.05) (**Fig. 2A**). Furthermore, while female *WT* mice experienced, on average, 0.22 embolic events per minute, the female *LPA* mice experienced significantly fewer embolic events (0.07 events/min; p < 0.05) (**Fig. 2B**). Similar results were found in the male mice: mean vessel occlusion time was shortened in the *LPA* mice compared to the *WT* mice (5.24 min versus 9.64 min; p < 0.05) (**Fig. 2C**) and the LPA mice experienced fewer embolic events than WT mice (0.06 min versus 0.20 min; p < 0.05) (**Fig. 2D**). Together, these findings demonstrate that the presence of apo(a) both accelerates thrombogenesis while also resulting in more stable clots.

**Fig. 2.**
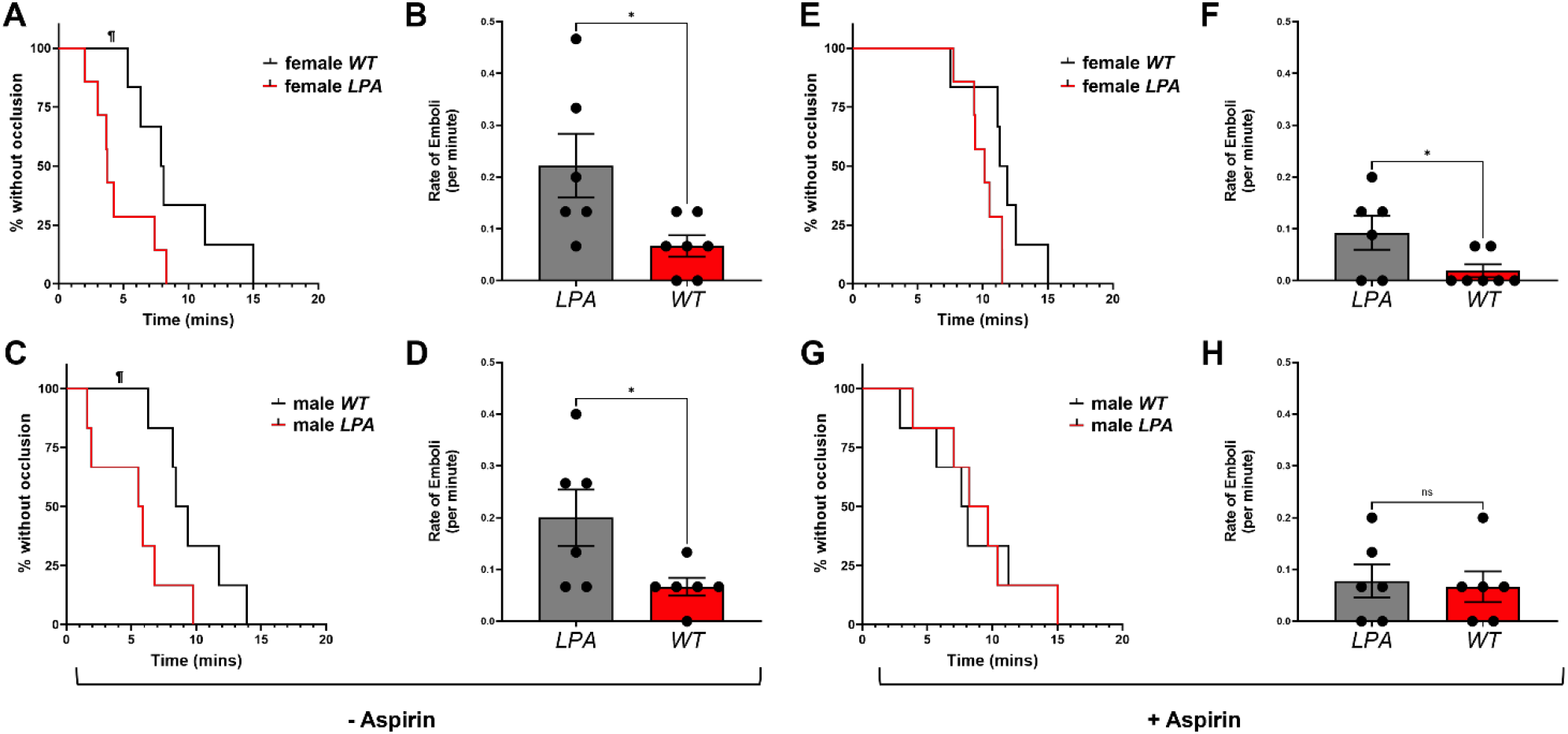
Apo(a) accelerates rates of vessel occlusion and decreases the number of embolic events in a FeCl_3_-induced carotid artery thrombosis model, with effects reversed by aspirin. (**A**) Kaplan-Meier curves depicting % of mice free from occlusive thrombi over time and (**B**) scatterplot showing distribution of rates of embolic events in wild-type (*WT*; n=6) and apo(a)-expressing (*LPA*; n=7) female mice. Scatterplots show mean of each group ± SEM. (**C**) Kaplan-Meier curves depicting % of mice free from occlusive thrombi over time and (**D**) scatterplot showing distribution of rates of embolic events in *WT* (n=6) and *LPA* (n=6) male mice. (**E**) Kaplan-Meier curves depicting % of mice free from occlusive thrombi over time and (**F**) scatterplot showing the distribution of rates of embolic events in wild-type (*WT*; n=6) and apo(a)-expressing (*LPA*; n=7) female mice receiving aspirin. (**G**) Kaplan-Meier curves depicting % of mice free from occlusive thrombi over time and (**H**) scatterplot showing distribution of rates of embolic events in *WT* (n=6) and *LPA* (n=6) male mice receiving aspirin. Scatterplots show the mean of each group ± SEM. All mice were fed a HFHC diet. †*p* < 0.05 by log-rank test; \**p* < 0.05 by Student’s t-test.

However, we found very different results in mice that had been administered aspirin. There was no significant difference in mean vessel occlusion times between *LPA* and *WT* mice in neither females nor males (**Fig. 2E,G**). Aspirin increased the occlusion time in *LPA* mice for both females (a 116% increase) and males (72.3% increase) (compare **Figs. 2A,C and 2E,G**), while having a much smaller effect (28.7% increase) in female *WT* mice and no effect in male *WT* mice. In other words, *LPA* mice received a benefit from aspirin (prolongation of occlusion time) that *WT* mice did not, an observation that is directionally consistent with the reported benefit of aspirin in high-Lp(a) patients in reducing MACE or cardiovascular death.^59^ While female *LPA* mice receiving aspirin still showed a reduced rate of embolic events compared to female *WT* mice receiving aspirin, this difference was abolished in male mice (**Fig. 2F,H**).

### Chandler loop thrombus analogues formed *in vitro* in the presence of Lp(a) are resistant to tPA-mediated fibrinolysis

Having demonstrated a direct prothrombotic and clot-stabilizing effect of apo(a) using two different thrombosis models in transgenic mice, we next sought to probe the underlying mechanisms involved. For these studies, we turned to the Chandler loop model, in which thrombi are formed from whole recalcified blood under flow conditions with a defined shear rate meant to mimic the hemodynamics of arteries susceptible to atherothrombosis.

We first assessed if Lp(a) and apo(a) impacted the lysis of the Chandler thrombi. We formed clots from whole blood supplemented with Alexa-Fluor-488-conjugated fibrinogen in the presence of vehicle control, or 250 nmol/L plasma-purified Lp(a) or 17K recombinant apo(a). We also tested two mutants of 17K: 17KΔLBS10 that lacks the strong lysine binding site in KIV10 which mediates interactions of apo(a) with biological substrates, and 17KΔP that lacks the protease domain which mediates binding to plasminogen and hence the antifibrinolytic effects of apo(a).^24,49^ After formation, thrombi were transferred to autologous plasma containing tPA and lysis was monitored by release of fluorescently labeled fibrin degradation products (flu-FDP) into the plasma. Compared to control clots, we observed robust and significant reductions of flu-FDP release when thrombi were formed in the presence of 17K wild-type apo(a) (up to 50%; **Fig. 3A**). Similarly, significant reductions in fibrinolysis were observed when clots were formed in the presence of either Lp(a) (up to 33.2%), 17KΔLBS10 (up to 35.5%) or 17ΔP (up to 37.8%), versus control.

**Fig. 3.**
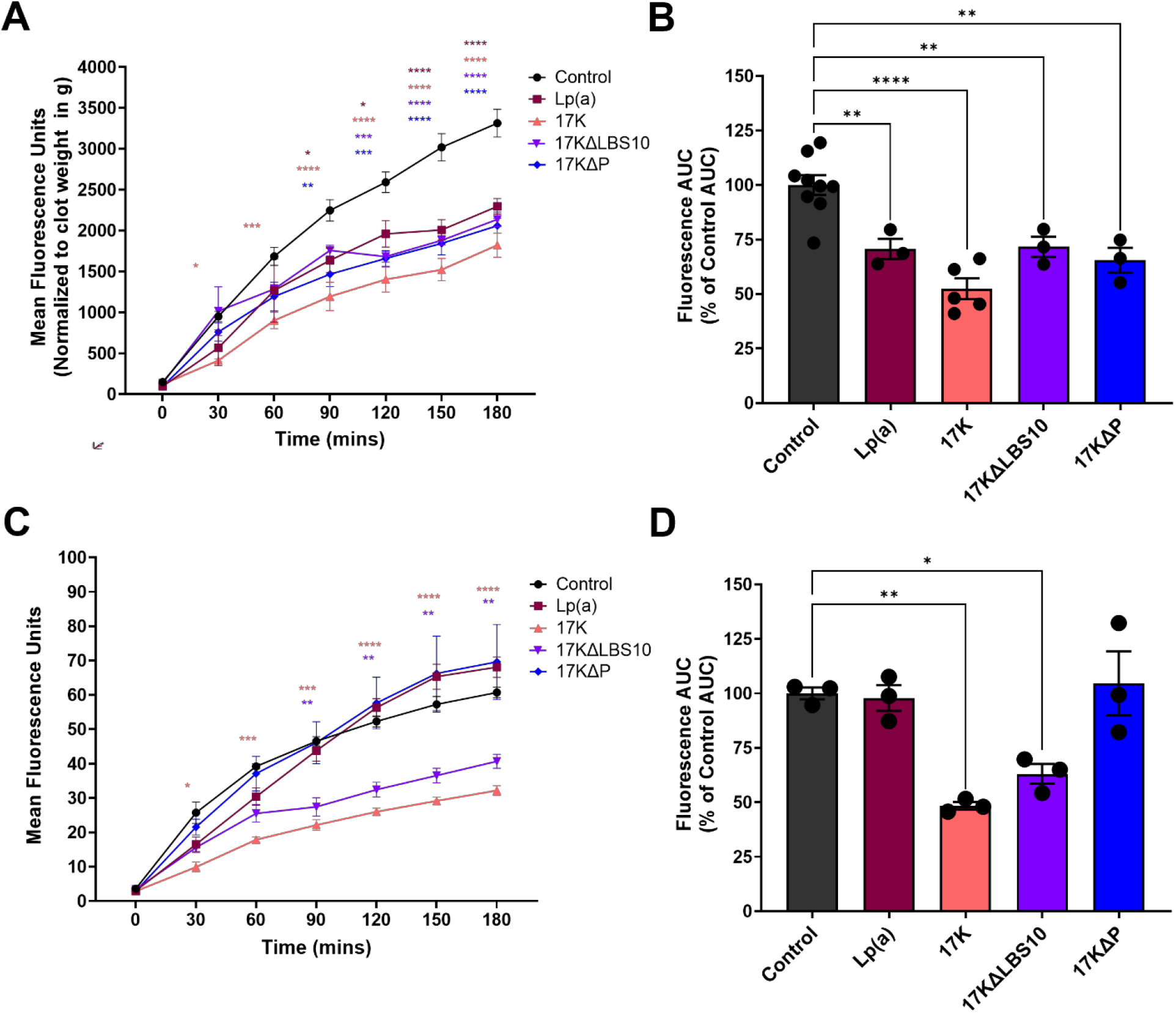
Effect of Lp(a) and apo(a) variants on fibrinolysis of clots formed from flowing whole blood in a Chandler loop apparatus. (**A**) Citrated whole blood collected from a human donor with low Lp(a) levels was supplemented with HEPES buffered saline (HBS; vehicle control) or 250 nmol/L of either plasma-purified Lp(a), 17K recombinant apo(a), 17KΔLBS10, or 17KΔP. The blood was recalcified and a thrombus was formed in a Chander loop. The thrombi were retrieved, washed, and bathed in autologous plasma containing tPA. Fibrinolysis was assessed as the release of fluorescent fibrin degradation products (flu-FDPs) from the clot into the plasma over time. (**B**) Chandler loop thrombi were formed (as described above) in the absence of added Lp(a) or apo(a). Thrombi were then bathed in autologous plasma containing tPA and either vehicle or 250 nmol/L Lp(a), 17K, 17KΔLBS10, or 17KΔP. Fibrinolysis was assessed as described above. Mean fluorescence readings were normalized to an averaged weight of clots formed in the presence of each variant. Left panels display Mean Fluorescence Unit readings measured every 30 minutes over a period of 180 minutes. Right panels display fluorescence data expressed as Area Under the Curve (AUC) and relative to AUC of control clots. Data represent mean ± SEM from at least 3 independent experiments. Significant differences compared to vehicle controls were determined using either two-way ANOVA (left panels) or one-way ANOVA (right panels) with Tukey multiple comparisons post-hoc analysis. \**p* < 0.05, \*\**p* < 0.01, \*\*\**p* < 0.001, \*\*\*\**p* < 0.0001.

It is worth noting that these experiments cannot resolve whether the observed decreases in flu-FDP release associated with the presence of Lp(a) or apo(a) were due to effects on platelet function or coagulation, or due to fibrinolysis inhibition. Pro-aggregatory (and/or pro-coagulant) effects on platelets or fibrin deposition and cross-linking, as well as thrombin-mediated antifibrinolytic effects via TAFI/CPB2 impacting the rate of plasminogen activation, would all presumably result in lysis-resistant thrombi and decreased release of flu-FDPs. To determine whether the above findings were due, in part, to direct antifibrinolytic effects, Chandler thrombi formed in the absence of Lp(a) or apo(a) were placed into autologous plasma containing tPA as well as either Lp(a), 17K, 17KΔLBS10, or 17KΔP.

Subsequently, fibrinolysis was determined by assessing the release of flu-FDPs into the plasma over time. In these experiments, any observable inhibitory effects of Lp(a) or apo(a) on rates of fibrinolysis would be on the surface of already formed clots, thus reflecting purely antifibrinolytic effects versus formation of lysis-resistant thrombus structures. We observed that compared to control, fibrinolysis was significantly decreased by exogenously added 17K (up to 47%; **Fig. 3B**) or 17KΔLBS10 (up to 36.2%). There were no significant differences in fibrinolysis compared to control from exogenously added Lp(a) or 17KΔP (**Fig. 3B**). These findings are consistent with our previous report demonstrating that while 17K and 17KΔLBS10 inhibit plasma clot lysis, Lp(a) and 17KΔP do not. They also indicate that the effect of Lp(a) on thrombus lysis would be attributable to Lp(a) promoting a lysis-resistant clot structure; by contrast, apo(a) can both promote a lysis-resistant clot structure and inhibit lysis directly.

### Lp(a) promotes platelet and PAI-1 accumulation during thrombogenesis of Chandler loop clot analogues

We sectioned and characterized by immunofluorescence Chandler thrombi formed in the presence of Lp(a), 17K, 17KΔLBS10, or 17KΔP to discern effects on platelet and fibrin deposition that might explain their lysis resistance. We observed that compared to control clots, CD42b (platelet marker GPIb) staining was significantly increased for thrombi formed in the presence of Lp(a), 17K, and 17KΔP (**Fig. 4A,B**); no differences in CD42b staining were observed between control thrombi or those formed in the presence of 17KΔLBS10 (**Fig. 4A,B**). Interestingly, using an anti-apo(a) antibody for co-immunofluorescence, Lp(a), 17K, and 17KΔP staining appeared to be abundantly present in regions rich in platelets, whereas 17KΔLBS10 did not appear to deposit as substantially in these regions (**Fig. 4A**). We also assessed the abundance of apo(a) staining relative to total clot area. Apo(a) was undetectable in the absence of added Lp(a) or apo(a) (**Fig. 4B**), and 17KΔLBS10 staining was less abundant than that of Lp(a), 17K, or 17KΔP (**Fig. 4C**).

**Fig. 4.**
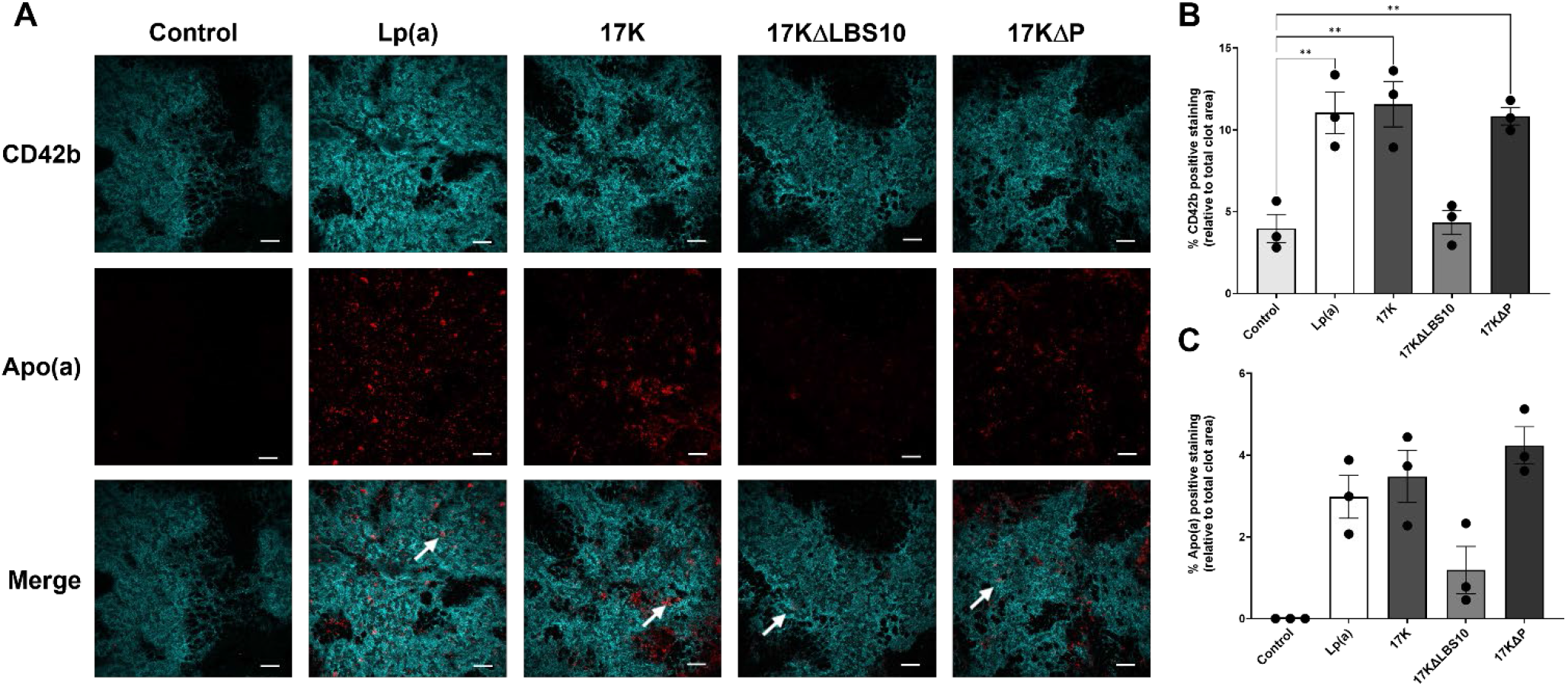
Effect of Lp(a) and apo(a) variants on platelet accumulation in Chandler loop clots. (**A**) Representative images of Chandler loop clot sections co-stained to detect the presence of apo(a) (red) and platelets (CD42b; cyan). Clots were formed in the presence of HBS (control) or 250 nmol/L Lp(a), 17K, 17KΔLBS10, or 17KΔP. Following thrombogenesis in the Chandler loop apparatus, clots were fixed and subsequently sectioned longitudinally in 5-µm steps prior to immunofluorescent staining. Z-stack images of clot sections were acquired under identical acquisition settings and excitation wavelengths using the Nikon AIR+ confocal laser scanning system equipped with a 60× oil-immersion objective and high-resolution Galvano scanner. Representative micrographs display z projections of z-stack image files (21 total z-slices, step size of 0.2 µm, 1× digital zoom). All images are displayed at the same brightness/intensity scale. Overlay images contain pseudocoloured cyan (488 nm emission) and colour composite red (647 nm emission). Clot areas and % positive staining for each emission wavelength were assessed using QuPath software. Quantitative data displaying fractional surface coverage (%) by (**B**) platelets (CD42b; cyan) or (**C**) apo(a) (red) relative to total clot areas is shown. Data represent mean ± SEM from 3 independent Chandler loop clot generation experiments. Significant differences compared to control clots (**B**) or to Lp(a)-containing clots (**C**) were determined using one-way ANOVA with Dunnett’s post-hoc analysis. \*\**p* < 0.01.

The main reservoir of the tPA inhibitor PAI-1 in circulation is located within platelet α-granules.^60^ Furthermore, it has been shown that a functional pool of platelet PAI-1 anchors to the surface of activated platelets and regulates fibrinolysis locally.^61^ Accordingly, we found that PAI-1 staining appeared to localize to regions that also stained positively for platelets **(Supplementary Fig. S5**). Consistent with our quantitative platelet positive staining findings, we observed that PAI-1 staining was significantly increased in thrombi formed in the presence of either Lp(a), 17K, or 17KΔP when compared to control clots (**Fig. 5**); no significant differences in PAI-1 staining were observed between control clots and clots formed in the presence of 17KΔLBS10. Qualitatively, within clots formed in the presence of either Lp(a), 17K, or 17KΔP, we observed an abundance of apo(a)-positive staining in regions rich in PAI-1 (**Fig. 5A**). Conversely, apo(a)-positive staining was less abundant in PAI-1-rich regions within the 17KΔLBS10 clots. These qualitative findings are consistent with what was observed for clot sections stained for platelets (**Fig. 4A**).

**Fig. 5.**
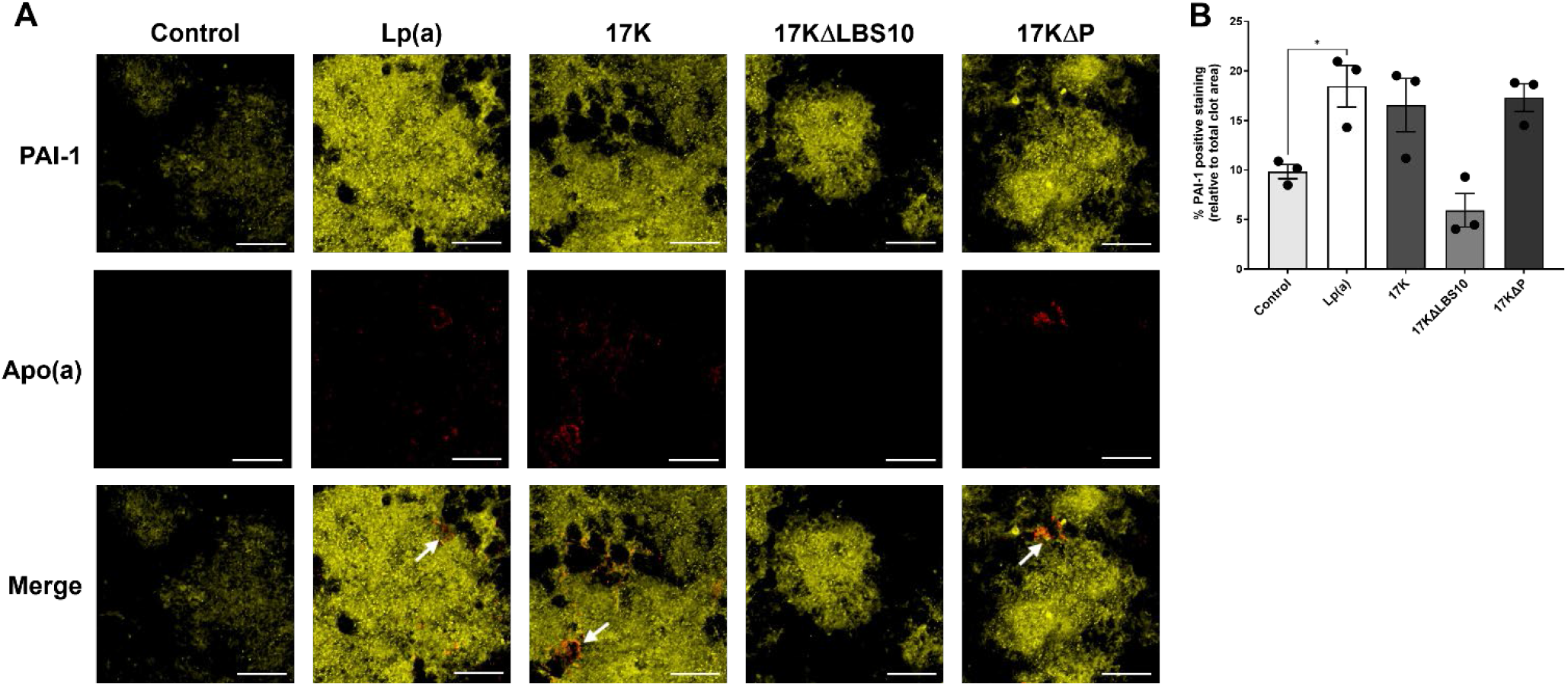
Effect of Lp(a) and apo(a) variants on PAI-1 accumulation in Chandler loop clots. (**A**) Representative images of Chandler loop clot sections co-stained to detect the presence of apo(a) (red) and plasminogen activator inhibitor-1 (PAI-1; yellow). Clots were formed in the presence of HBS (control) or 250 nmol/L Lp(a), 17K, 17KΔLBS10, or 17KΔP. Following thrombogenesis in the Chandler loop apparatus, clots were fixed and subsequently sectioned longitudinally at 5-µm steps prior to immunofluorescent staining. Z-stack images of clot sections were acquired under identical acquisition settings and emission wavelengths using the Nikon AIR+ confocal laser scanning system equipped with a 40× oil-immersion objective and high-speed Resonant scanner. Representative micrographs display z projections of z-stack image files (13 total z-slices, step size of 0.925 µm, 3× digital zoom). Images are displayed at the same brightness/intensity scale. Overlay images are pseudocoloured yellow (647 nm emission) and red (488 nm emission). White arrows indicate regions of apparent co-localization of apo(a) positive staining with PAI-1 positive staining. Scale bars = 20 µm. (**B**) Quantitative data displaying fractional surface coverage (%) by PAI-1 (yellow) relative to total clot areas. Data represent mean ± SEM from at least 3 independent Chandler loop clot generation experiments. Significant differences compared to vehicle control clots were determined using one-way ANOVA with Dunnett’s post-hoc analysis. \**p* < 0.05.

### Chandler loop thrombus analogues formed in the presence of Lp(a) contain fibrin networks with features of fibrinolysis resistance

The density of fibrin staining was determined by quantifying the surface area of green fluorescence in randomly selected confocal images of fibrin-positive staining regions (**Fig. 6A**); as expected, fibrin was the predominant structure observed in all Chandler loop thrombi (**Supplementary Fig. S5**). Compared to control, we observed that fibrin densities were significantly increased within clots formed in the presence of Lp(a), 17K, and 17KΔP, but not 17KΔLBS10 (**Fig. 6B**). Fibrin network porosity was also manually determined from the same confocal images: decreased porosities and increased fibrin fiber densities, which can be attributed to either increased fibrin deposition and cross-linking or fibrinolysis inhibition, have been shown to preclude appropriate clot breakdown.^62–64^ We observed that pore diameters were significantly decreased in Lp(a), 17K, and 17KΔP thrombi when compared to control clots (**Fig. 6C**), with no differences in pore diameters observed between control clots and 17KΔLBS10 clots (**Fig. 6C**).

**Fig. 6.**
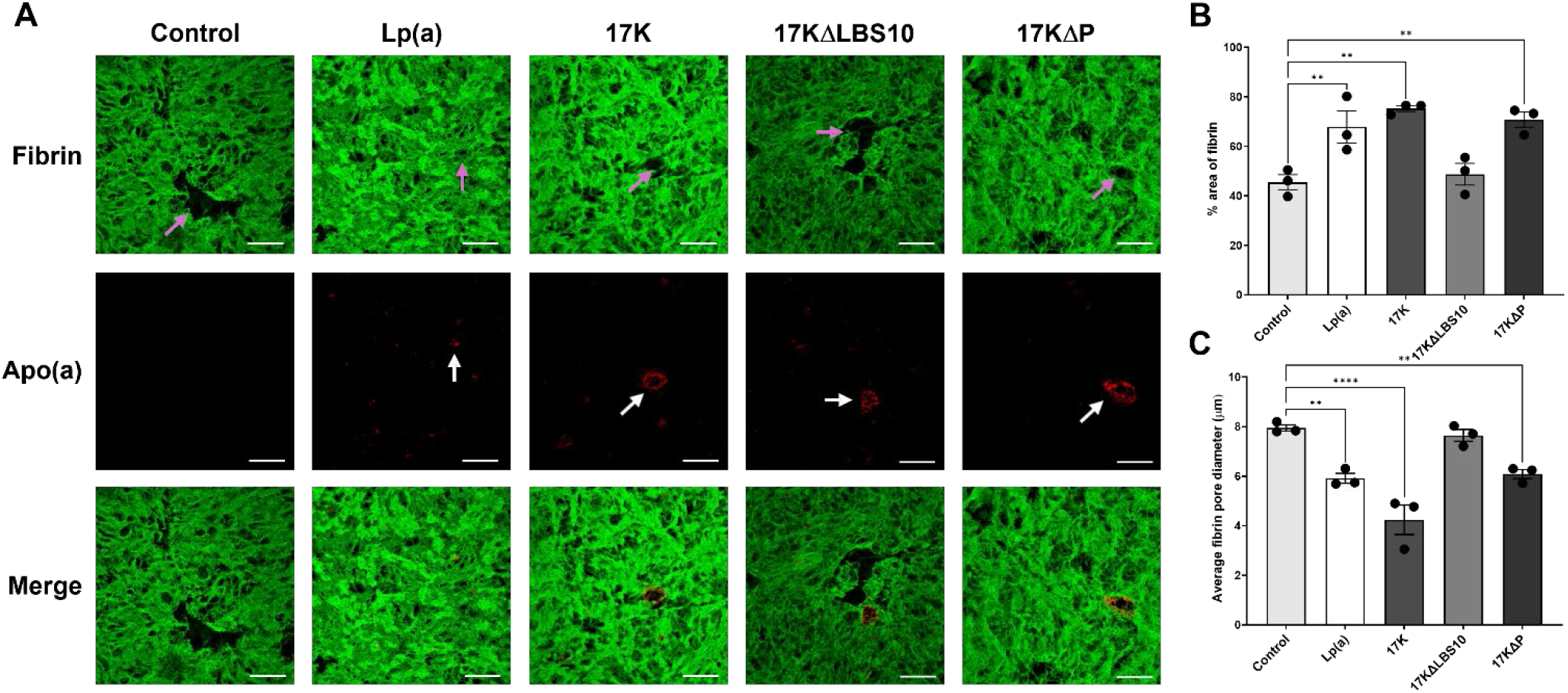
Fibrin network densities and pore diameters of Chandler loop clots formed in the presence of Lp(a) or apo(a) variants. (**A**) Representative images of Chandler loop clot sections co-stained to detect the presence of apo(a) (red) and fibrin (green). Clots were formed in the presence of HBS (control) or 250 nmol/L Lp(a), 17K, 17KΔLBS10, or 17KΔP. Following thrombogenesis in the Chandler loop apparatus, clots were fixed and subsequently sectioned longitudinally in 5-µm steps prior to immunofluorescent staining. Z-stack images of clot sections were acquired under identical acquisition settings and emission wavelengths using the Nikon AIR+ confocal laser scanning system equipped with a 40× oil-immersion objective and high-speed Resonant scanner. Representative micrographs display z projections of z-stack image files (13 total z-slices, step size of 0.925 µm, 3× digital zoom). Images are displayed at the same brightness/intensity scale. Overlay images are pseudocoloured green (488 nm emission) and red (647 nm emission). Pink arrows indicate fibrin pores. White arrows indicate distinctive circular structures of Lp(a) and apo(a) within the fibrin-rich regions. Scale bars = 20 µm. (**B**) Quantitative data displaying mean fractional surface coverage (%) by fibrin (green). Fractional surface coverage by fibrin was used as a measure of fibrin density. For each replicate (n=3), six 40× magnification images were randomly acquired throughout the fibrin-rich tail region of the clots. Density measurements were averaged across the 6 random images to determine an average fibrin density for each replicate. (**C**) Average pore diameters were determined using the same 40× magnification images described above. Data represent mean ± SEM from 3 independent Chandler loop clot generation experiments. Significant differences compared to control clots were determined using one-way ANOVA with Dunnett’s post-hoc analysis. \*\**p* < 0.01, \*\*\*\**p* <0.0001.

Notably, apo(a)-positive staining was consistently visible as unique circular patterns within clots containing Lp(a) or apo(a) (**Fig. 6A**). Activated coagulation Factor XIII (FXIIIa) influences clot stabilization through catalyzing the covalent cross-linking of ε-(γ-glutamyl)lysine isopeptide bonds between fibrin α- or γ-chains.^65^ The presence of FXIIIa during clot formation has also been associated with increased densities of fibrin fiber deposition and reduced pore sizes.^66^ Moreover, FXIIIa has also been shown to facilitate fibrinolysis inhibition through intermolecular cross-linking of other proteins to fibrin, such as α2-antiplasmin.^67^ Interestingly, it has previously been shown that FXIIIa is also capable of cross-linking Lp(a) to fibrin within human atherosclerotic lesions.^68^ In our Chandler loop thrombi, we observed that the patterns of circular apo(a) immunofluorescence staining within Lp(a) and apo(a) clots tended to preferentially localize to regions that also stained positively for FXIIIa (**Supplementary Fig. S6**).

### SEM analysis of Lp(a) and apo(a)-induced changes to the ultrastructural composition of thrombi formed in the Chandler loop apparatus

SEM was used to quantitatively assess the impact of Lp(a) and apo(a) on changes to the ultrastructural composition of thrombi formed in the Chandler loop apparatus (**Fig. 7**). While red blood cells (RBCs) were the predominant cell type observed in thrombi formed in the presence of either vehicle control, Lp(a) or 17K apo(a), RBC content was significantly decreased in clots formed in the presence of Lp(a) or in the presence of 17K (**Fig. 7G**). More strikingly, RBC shape changes were evident when comparing vehicle control clots with Lp(a) or apo(a) clots (**Fig. 7A-C**). In control clots, the vast majority of RBCs had the typical biconcave appearance (**Fig. 7H,I**). However, in both the Lp(a) and 17K clots, almost all the RBCs were polyhedrocytes (**Fig. 7J**). This form of RBCs, first reported by the Weisel group, represents tightly packed RBCs that have been deformed by clot contraction into polyhedral shapes that occupy the minimum possible volume.^33^

**Fig. 7.**
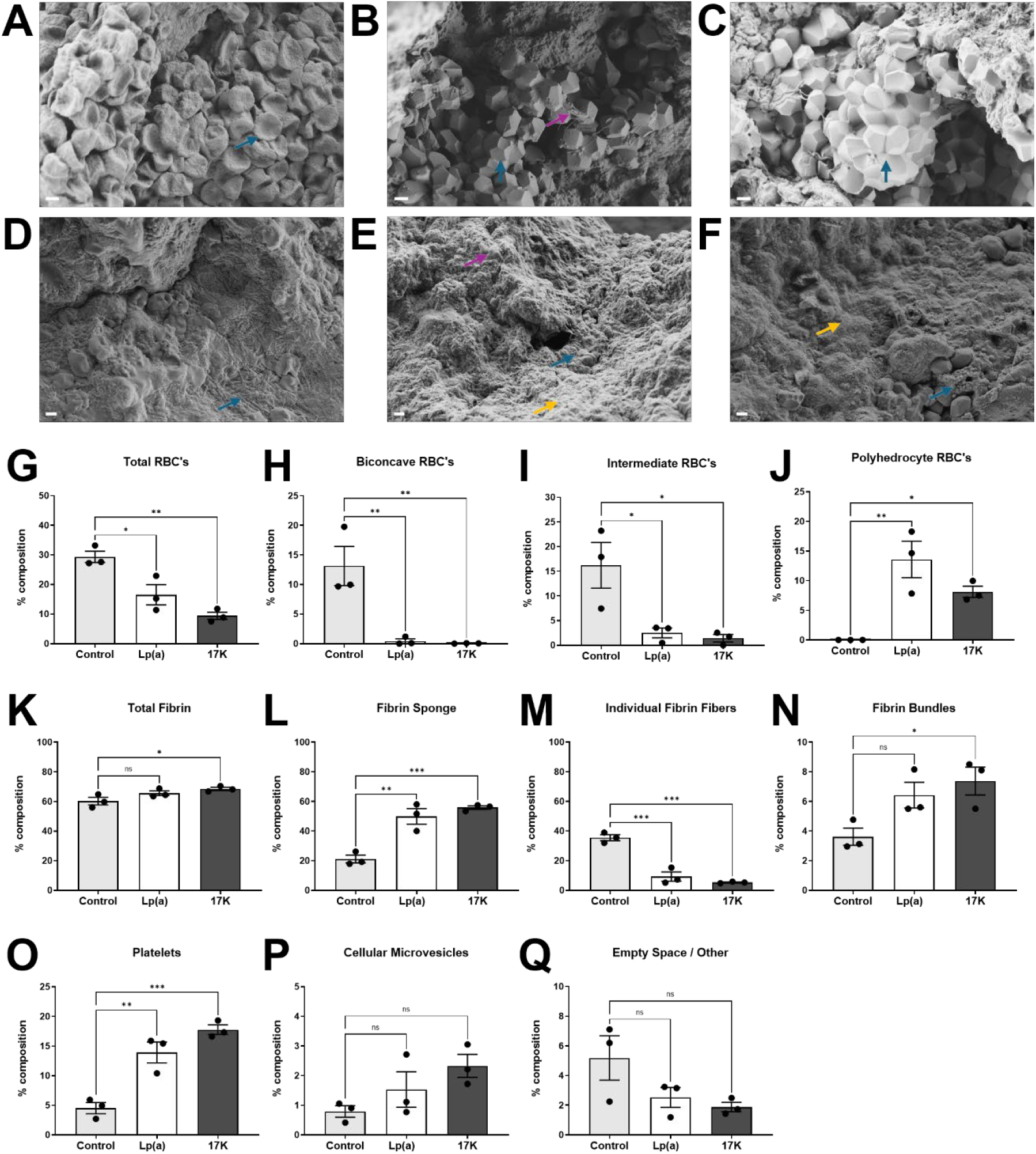
Effects of Lp(a) and apo(a) variants on Chandler loop clot ultrastructures assessed using scanning electron microscopy. Representative scanning electron microscopy (SEM) micrographs of red blood cell (RBC)-rich regions (**A-C**) and fibrin-rich regions (**D-F**) of Chandler loop thrombi. (**A**) RBC morphology of vehicle control clots, where the blue arrow indicates biconcave RBC. (**B,C**) RBC morphology of Lp(a) and 17K clots, respectively. The pink arrow indicates fibrin bundles; blue arrows indicate polyhedrocyte RBCs. (**D**) Fibrin-rich region of vehicle control clots, where the blue arrow indicates individual fibrin fibers. (**E**) Fibrin-rich region of Lp(a) clots, where the orange arrow indicates fibrin sponge, the blue arrow indicates individual platelet with cell surface projections, and the pink arrow indicates platelet aggregate. (**F**) Fibrin-rich region of 17K clot, where the orange arrow indicates fibrin sponge and the blue arrow indicates cellular microvesicles. Scale bars = 2 µm. (**G-Q**) Quantitative analyses of structures (composition %) identified within SEM images of Chandler loop thrombi formed in the presence of either vehicle control, Lp(a) or 17K. Structures were identified and quantified as described in Materials and Methods. All SEM images used for analysis were acquired at 2000× magnification. Data represent mean ± SEM from 3 independent Chandler loop clot generation experiments. Significant differences compared to control were assessed statistically by one-way ANOVA with Tukey post-hoc analysis. \**p* < 0.05, \*\**p* < 0.01, \*\*\**p* < 0.001, ns = not significant.

Within clots formed in the presence of either vehicle control, Lp(a), or 17K apo(a), fibrin structures presented as individual fibers, fibrin bundles, and fibrin sponge (**Fig. 7D-F**). While total fibrin content did not differ between control clots and Lp(a) clots, thrombi formed in the presence of 17K had significantly increased amounts of fibrin deposition compared to control (**Fig. 7K**). Although only marginal changes were observed in total fibrin content, more robust differences were observed when comparing the proportions of each individual fibrin structure. Fibrin sponge content was significantly increased in Lp(a) and 17K clots compared to control (**Fig. 7L**). Conversely, individual fibrin fiber content was significantly decreased in Lp(a) and 17K clots compared to control clots (**Fig. 7M**). Fibrin bundle content was significantly increased in 17K clots compared to control, with an upward trend in Lp(a) clots (**Fig. 7N**). Platelets accounted for 13.9% of total clot content in Lp(a) clots and 17.8% in 17K clots; this was significantly increased compared to control clots (4.5%; **Fig. 7O**). While no statistical differences were observed, cellular microvesicle content trended upward in Lp(a) and 17K clots compared to control (**Fig. 7P**). Furthermore, the proportion of empty space compared to control trended downwards in the Lp(a) and 17K clots (**Fig. 7Q**).

### Lp(a) levels and arterial or venous thrombosis in participants of the UK Biobank

The baseline characteristics of participants included in the arterial and venous thrombosis analyses, stratified by whether they experienced either arterial embolism or thrombosis (as defined by IDC10 code I74; see Material and Methods) or venous thromboembolism (VTE), or neither event during follow-up are presented in **Supplementary Table S1**. Among all participants, 73,483 (16.5%) had an elevated Lp(a) level (≥125 nmol/L). Of the participants in the “low Lp(a)” group, 1035 had an incidence of arterial embolism or thrombosis and 8818 had an incidence of VTE. Those with Lp(a) ≥125 nmol/L were at significantly increased risk for arterial embolism or thrombosis (hazard ratio (HR) 1.38 (95% confidence interval (CI) 1.21-1.57; p = 1.28 ×10^6^) (**Fig. 8A**). The increased risk was significant for men, but not for women, and was not affected by age, being similar in those above or below the median age (58 years) (**Supplementary Table S2**). By contrast, Lp(a) ≥125 nmol/L did not increase risk for VTE (HR (95% CI) 1.02 (0.97-1.08); p = 0.366) (**Fig. 8B**), which held true for both men and women and both age categories (**Supplementary Table S2**).

**Fig. 8.**
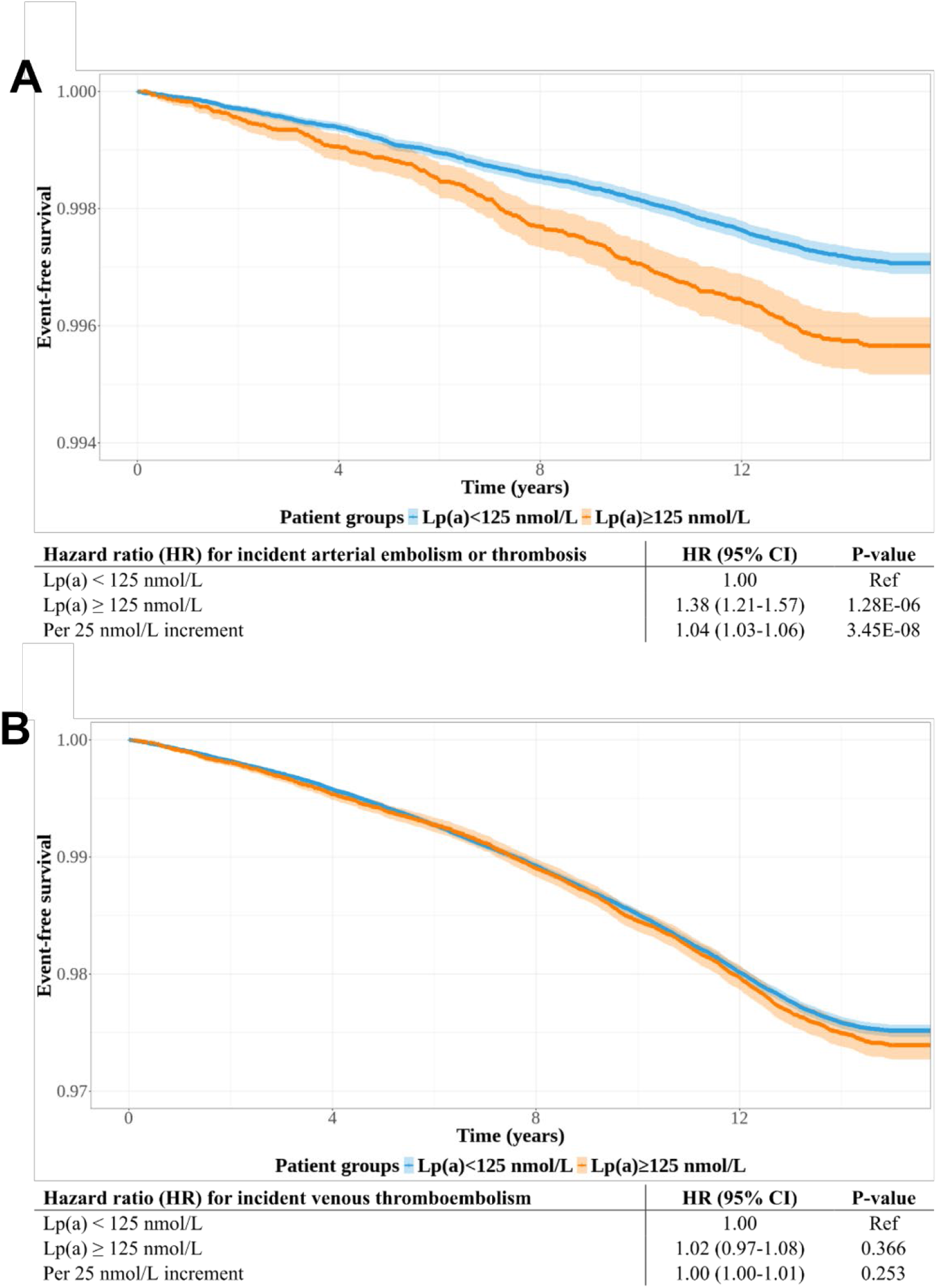
Elevated Lp(a) is a risk factor for arterial thrombosis but not venous thrombosis in UK Biobank participants. (**A**) Kaplan-Meier plot of the event-free survival for first incidence of arterial thrombosis, defined using the ICD-10 code I74, which denotes the incidence of arterial embolism or thrombosis. Hazard ratios for event-free survival with high Lp(a) (≥125 nmol/L) versus low Lp(a) (<125 nmol/L) as well as per 25 nmol/L increase in Lp(a) for the incidence of arterial embolism or thrombosis are indicated. (**B**) Kaplan-Meier plot of the event-free survival for first incidence of VTE, a composite of deep vein thrombosis, pulmonary embolism, or venous thrombophlebitis (ICD-10 I80.2, I82.2, I26.0, I26.9, OPCS-4 codes L79.1 and L90.2). Hazard ratios for event-free survival with high Lp(a) (≥125 nmol/L) versus low Lp(a) (<125 nmol/L) as well as per 25 nmol/L increase in Lp(a) for the incidence of VTE are indicated.

## DISCUSSION

In this work, for the first time, we have demonstrated a prothrombotic role for apo(a) in two distinct thrombosis models *in vivo*. Moreover, we show for the first time that Lp(a) (and apo(a)) delays thrombolysis of whole blood clots formed under arterial flow conditions. Characterization of these thrombus analogues by immunofluorescence provided evidence for increased platelet deposition and PAI-1 release, increased fibrin density, and reduced fibrin porosity attributable to Lp(a) and apo(a). Most strikingly, SEM images of thrombi formed in the presence of Lp(a) and apo(a) show a dramatic alteration of clot ultrastructure, with increased fibrin sponge, a predominance of fibrin bundles versus individual fibers, and evidence of substantially increased clot contraction as revealed by the deformation of erythrocytes into polyhedrocytes. In a novel analysis of UK Biobank data, we found that elevated Lp(a) increased risk for arterial thrombosis and embolism, which is identified in its ICD-10 code as embolism and thrombosis of the aorta, upper and lower extremities, iliac artery, and other unspecified arteries. Importantly, this ICD-10 code does not include coronary, carotid, and cerebral embolism and thrombosis, encompassing acute myocardial infarction and ischemic stroke, as well as intermittent claudication due to peripheral artery disease. Therefore, we have identified a direct role for Lp(a) in arterial embolism or thrombosis, independent of its recognized contributions to atherosclerosis processes. Together with our *in vitro* and animal model thrombosis data, these findings constitute compelling evidence that Lp(a) is an inherently prothrombotic lipoprotein.

The role of Lp(a) as a prothrombotic factor is not well understood and constitutes a highly understudied and controversial area of Lp(a) pathophysiology. Since the discovery that apo(a) is homologous to the fibrinolytic proenzyme plasminogen, this question has been dominated by assumption that Lp(a) has the ability to interfere with fibrinolysis.^21,69^ However, recent studies from our group have cast doubt on this assumption, since Lp(a) lowering does not affect *ex vivo* plasma clot lysis times and the presence of the lipoprotein particle in Lp(a) obscures the antifibrinolytic potential of apo(a).^23,24^ However, we also found evidence that Lp(a) and apo(a) can stimulate coagulation and thrombin generation in plasma^24^; together with earlier reports that Lp(a) and apo(a) can potentiate platelet aggregation in response to different agonists^34–36^, this motivated us to examine thrombosis in our transgenic mice expressing high levels of apo(a).

In the present study, we show that compared to wild-type mice, mice expressing apo(a) have significantly enhanced platelet and fibrin volumes in a laser-induced mesenteric vessel thrombosis model, particularly in the apo(a) mice fed a HFHC diet. Existing *in vitro*/*ex vivo* evidence is conflicting regarding the effect of Lp(a) on platelet reactivity. Specifically, the direction of this relationship appears to be dependent on the particular platelet activating factors present.^34,35,70–73^ Several studies in human patient cohorts have failed to detect an association between levels of Lp(a) (or of OxPL-apoB, which are highly correlated to Lp(a)), and platelet reactivity in response to a variety of agonists.^72,74,75^ Importantly, however, the impact of Lp(a)/apo(a) on platelet activity has never been assessed within an *in vivo* context, where numerous pro-aggregatory factors simultaneously drive platelet biology, or under conditions of flow including *ex vivo* models. Taken together, our findings strongly suggest that Lp(a) and apo(a) exert prothrombotic effects through platelets. In addition to the increased platelet volumes observed in our intravital microscopy experiments, we found that administration of aspirin abolished the increased rate of occlusive thrombosis in mice expressing apo(a) in our FeCl_3_-induced carotid injury model. Moreover, we found that inclusion of Lp(a) or apo(a) in whole blood thrombi formed at arterial shear rates *ex vivo* enhanced the deposition of both platelets and PAI-1 in the thrombi.

The increased fibrin volumes in our apo(a)-expressing mice could be a consequence of increased platelet activation/aggregation, stimulation of the coagulation cascade (including through platelet-mediated mechanisms), or impaired fibrinolysis. The apo(a) component of Lp(a) has been observed to display dose-dependent antifibrinolytic effects in *in vitro*, *ex vivo*, and *in vivo* studies.^22–24,76^ However, the largely constant thrombus fibrin volumes observed during the thrombosis time course in both apo(a)-expressing and control mice are suggestive of limited net fibrinolytic activity, and certainly do not point to a difference in fibrinolytic activity between apo(a)-expressing and control mice.

Interestingly, administration of the thrombin inhibitor dabigatran in mice undergoing intravital imaging of laser-induced thrombosis was reported to be associated with dose-dependent reductions in fibrin volumes in developing thrombi.^45^ Taken together, it appears that the increased fibrin volumes in the apo(a) mice are related to the effect of apo(a) on platelet activation and thrombin generation. This idea is strengthened by our findings in the Chandler loop model, in which deletion of the apo(a) protease domain (17KΔP), which is responsible for mediating apo(a)’s antifibrinolytic effects^24^ did not impact the ability of apo(a) to promote fibrin deposition. However, Lp(a) and apo(a) do appear to contribute, albeit indirectly, to lysis resistance of whole-blood thrombi: our data indicate that the enhanced platelet responses result in higher levels of PAI-1 within the thrombi as well as increased clot contraction, and the stronger procoagulant stimulus gives rise to lysis-resistant fibrin structures such as fibrin sponge.

Intriguingly, our experiments using recombinant variants of apo(a) offer important mechanistic insights. First, all effects of Lp(a) in our Chandler loop experiments were recapitulated by apo(a), indicating that it is the apo(a) moiety of Lp(a) that is mediating the prothrombotic effects. Second, the 17KΔP variant that cannot bind to plasminogen and cannot inhibit fibrinolysis^24,49^ still, like Lp(a), promoted a lysis-resistant thrombus phenotype when incorporated into the developing Chandler loop clots. Therefore, the lysis-resistant phenotype of thrombi observed in our *ex vivo* and *in vivo* experiments appears to be mediated by the effects of Lp(a) on platelets and coagulation. Third, the 17KΔLBS10 variant, which lacks the strong lysine-binding site in KIV10 as well as the covalently attached OxPL in this kringle^77^, was alone in failing to increase platelet and PAI-1 deposition or to stimulate denser and less porous fibrin formation. While this variant did inhibit thrombolysis of whole-blood clots, we attribute this to the antifibrinolytic ability of this variant, which our current data and previous studies in plasma^23^ show is equivalent to that of 17K itself. Intriguingly, however, numerous studies have shown a key role for platelet CD36 in mediating enhanced platelet responses in the context of oxidative stress.^78^ One key ligand for CD36 is OxPL, including the OxPL that decorate Lp(a) and are linked to apo(a) via KIV10.^13,14^ Therefore, it is tempting to speculate that the proplatelet and procoagulant effects of Lp(a) and apo(a) could be mediated by OxPL.

A notable limitation of our transgenic mouse thrombosis studies is that the mice express human apo(a), but not human apoB. Therefore, *bona fide* covalent Lp(a) particles are not formed in our animals since mouse apoB lacks the unpaired cysteine required for covalent linkage with apo(a), although non-covalent interactions are preserved.^79^ This raises the possibility that our prothrombotic effects are due to the presence of free apo(a) and perhaps would not be recapitulated by Lp(a). Several lines of evidence argue against this, however. First, our *ex vivo* studies clearly show that all prothrombotic effects of apo(a), except direct inhibition of fibrinolysis, are similarly evoked by Lp(a). Second, the lack of differences in apparent fibrinolysis in our mesenteric vessel laser-induced thrombi as well as the ability of aspirin to ablate the effect of apo(a) on increasing the rate of occlusion in the FeCl_3_-mediated carotid injury model indicate that it is enhanced platelet activation and coagulation rather than inhibition of fibrinolysis that accounts for the prothrombotic effect of apo(a) in the mice. Third, size exclusion chromatography of plasma from HFHC-fed apo(a)-expressing mice shows that apo(a) elutes from the column in the VLDL and LDL size ranges but not in later fractions even well beyond where HDL elutes (**Supplementary Fig. S7**). A similar finding was made in mice expressing the same *LPA* transgene in the context of *Ldlr* knockout and fed a standard diet.^42^ Hence, apo(a) is present as non-covalent complexes with mouse apoB-containing lipoproteins which we would predict – as for covalent Lp(a) particles – impairs the antifibrinolytic effects of apo(a).

If, as our data indicate, Lp(a) can exacerbate thrombosis, why is it that elevated Lp(a) does not increase risk for VTE? A key element of the pathogenesis of venous thrombosis is blood stasis, which is not a feature of either of our *in vivo* thrombosis models or of the *ex vivo* Chandler loop approach. Accordingly, platelets play a lesser role in the formation of venous thrombi which are also are less platelet-rich than arterial thrombi.^32,33^ Therefore, if apo(a) does in fact promote thrombosis via effects on platelet aggregation, which is consistent with our findings, these effects would likely only be pertinent to developing thrombi that are rich in platelets. We observed that apo(a) promoted thrombosis in both arteries (carotid) and veins (mesenteric) in response to distinct modalities of vascular damage, which would all presumably lead to platelet driven responses to the injury, analogous to arterial thrombosis. Nonetheless, extreme elevations of Lp(a) do appear to confer risk for VTE^29,31^, suggesting that Lp(a) promotes venous thrombosis, albeit less potently than it does arterial thrombosis. Another possibility is that arterial thrombosis involves a second “hit” by Lp(a), namely promotion of vulnerable atherosclerotic plaque features, that lowers the threshold for observation of an effect of Lp(a) on thrombosis.

In conclusion, our highly novel findings provide strong evidence that Lp(a) is inherently prothrombotic. This implies that the association of elevated Lp(a) with atherothrombotic disorders is mediated not only by the ability of Lp(a) to exacerbate atherosclerosis, but by the ability of this unique lipoprotein to enhance platelet activation and coagulation cascade activity. These findings align with the known contribution of elevated Lp(a) to non-atherosclerotic arterial thrombosis, including its association with ischemic stroke in children^80^ and with left atrial thrombus formation^41^, that is now extended by our novel analysis of UK Biobank data showing association of elevated Lp(a) with non-atherosclerotic arterial embolism and thrombosis events. Our results provide a mechanistic explanation for emerging data that aspirin use in a primary prevention setting reduces atherothrombotic events only in patients with elevated Lp(a). Moreover, our findings are relevant to Lp(a)-lowering therapies currently in clinical trials as they may have antithrombotic effects that would be realized much more rapidly than effects on the underlying atherosclerosis.

## ACKNOWLEDGEMENTS

The authors wish to thank Drs. Willam Lagor and Marcel Chuecos (Baylor College of Medicine) for their assistance in characterizing our transgenic mice expressing apo(a). We also thank Stephanie Milkovoch and the VITAL Facility at Robart Research Institute for their assistance in setting up the laser injury/intravital microscopy procedure and Dr. Filip Konecny for his assistance with performing the carotid artery thrombosis experiments.

## AUTHOR CONTRIBUTIONS

J.R.C., F.S.S., J.M.A., A.G., and S.T. performed the research; J.R.C., B.J.A., M.L.K., and M.B.B. designed the research; J.R.C. and M.B.B. drafted the manuscript, and all authors edited the manuscript.

