## Supplementary Information for "Lipoprotein(a) promotes thrombosis through platelet activation and promotion of a lysis-resistant thrombus architecture"

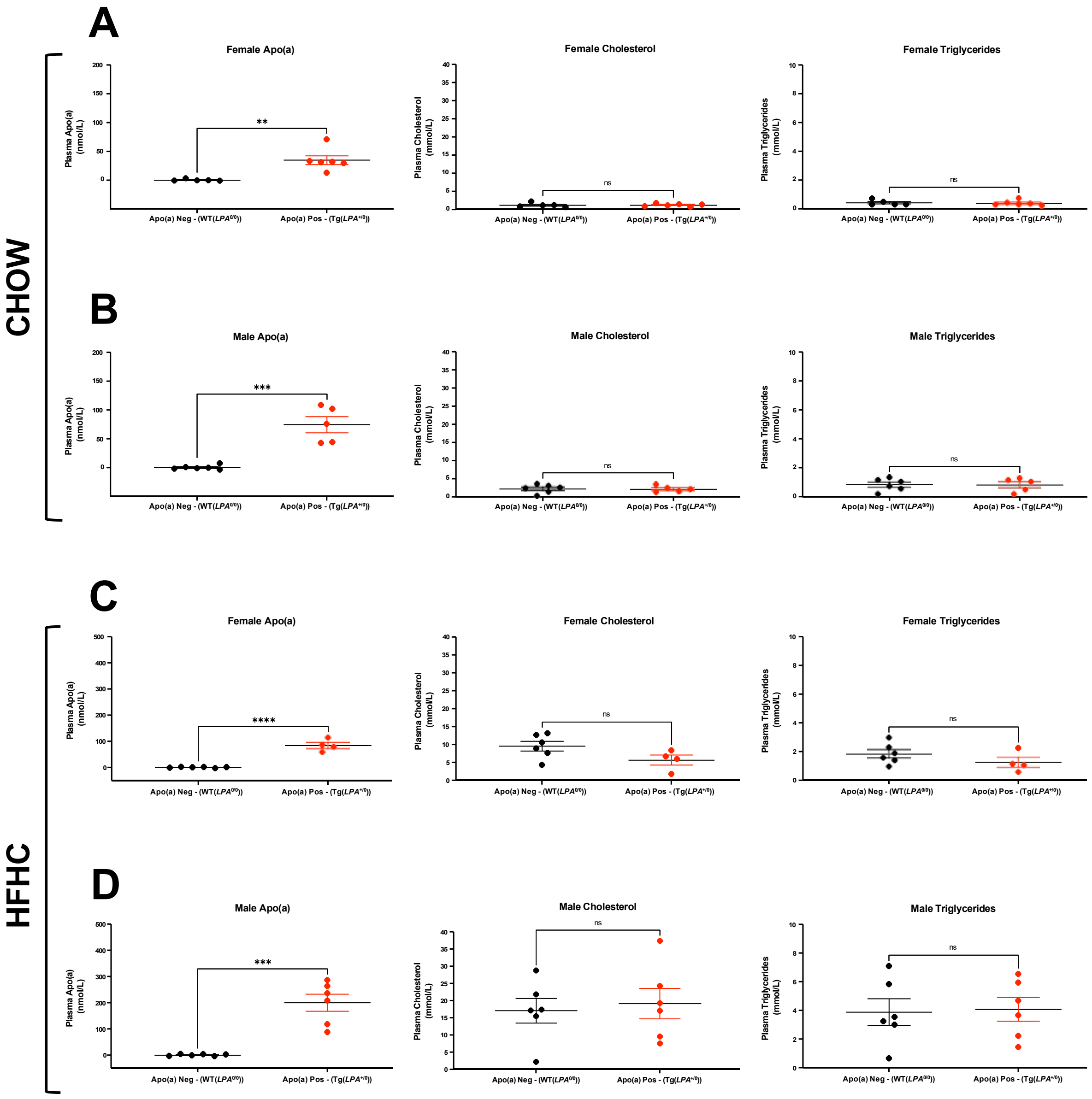


**Supplementary Fig. S1. Plasma apo(a), total cholesterol and triglyceride concentrations in wild-type (*WT*) and transgenic apo(a) (*LPA*) mice subjected to thrombosis studies**. Plasma apo(a), total cholesterol, and triglyceride levels were determined using Randox kits for (**A**) *WT* (n=5) and *LPA* (n=6) female mice fed chow diets; (**B**) *WT* (n=6) and *LPA* (n=5) male mice fed chow diets; (**C**) *WT* (n=6) and *LPA* (n=4) female mice fed high fat/high cholesterol (HFHC) diets; and (**D**) *WT* (n=6) and *LPA* (n=6) male mice fed HFHC diets. Data are means ± SEM of each group. Significant differences compared to *WT* mice animals were determined using Student's t-test. **$p < 0.01$, ***$p < 0.001$, ****$p < 0.0001$.

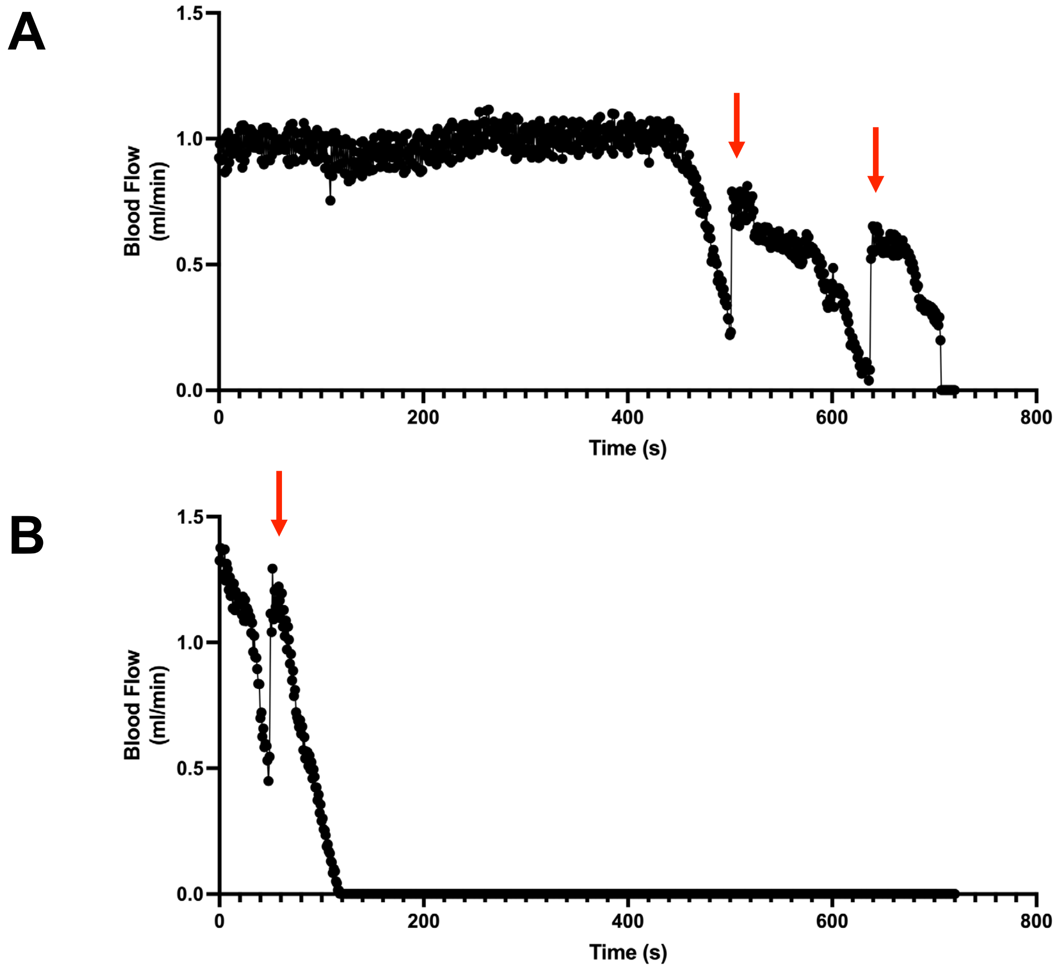


**Supplementary Fig. S2. Representative raw Doppler read-outs showing embolic events and cessation of blood flow in a carotid artery thrombosis model**. (**A**) *WT* and (**B**) *LPA* mice underwent $FeCl_3$-induced endothelial injury. Blood flow was monitored using a miniature Doppler flow probe (0.5PSB, Transonic System Inc.). Line-graphs show raw carotid artery Doppler blood flow readings over time in minutes (ml/min). Representative data are from male mice that did not receive low-dose aspirin therapy. Time 0 seconds (s) represents the baseline flow reading at the end of the $FeCl_3$ injury induction period. Average baseline blood flow rates within the RCA were 1.10 ml/min for male mice and 1.08 ml/min for female mice (not shown). Blood flow was continuously monitored using the flow probe from the time of baseline flow measurement, through the injury period, and up until a maximum of 15 minutes following the injury. The endpoints for the experiment were (i) when blood flow decreased to 25% of baseline flow (denoting 100% occlusion) or (ii) if occlusion was not observed within 15 minutes after injury. For the latter, 15 minutes was used as the value for statistical analyses. The number of embolic events observed per 15-minute Doppler reading was also recorded. An embolic event was identified as a sudden spike in blood flow during a period of general flow decline (red arrows).

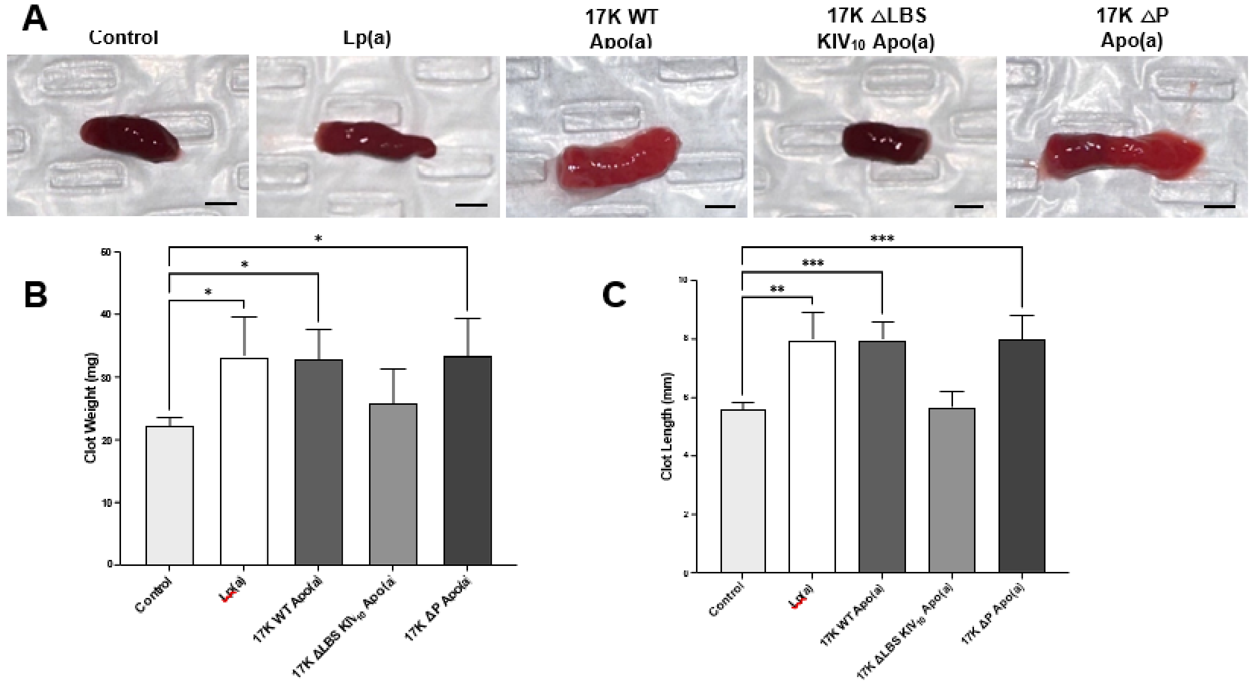


**Supplementary Fig. S3. Assessment of Chandler loop clot gross morphologies**. (**A**) Representative macroscopic images, (**B**) clot weights, and (**C**) clot lengths of Chandler loop thrombi formed in the presence of HBS (control) or 250 nmol/L Lp(a), 17K apo(a), 17KΔLBS10, or 17KΔP. Data represent means ± SEM from at least 4 independent experiments. Significant differences compared to vehicle control were determined using one-way ANOVA with Tukey post-hoc analysis. *$p < 0.05$, **$p < 0.01$, ***$p < 0.001$. Scale bars = 2 mm. Note that the clots containing Lp(a), 17K, and 17KΔP are lighter in colour and significantly greater in mass and length compared to control clots or clots containing 17KΔLBS10.

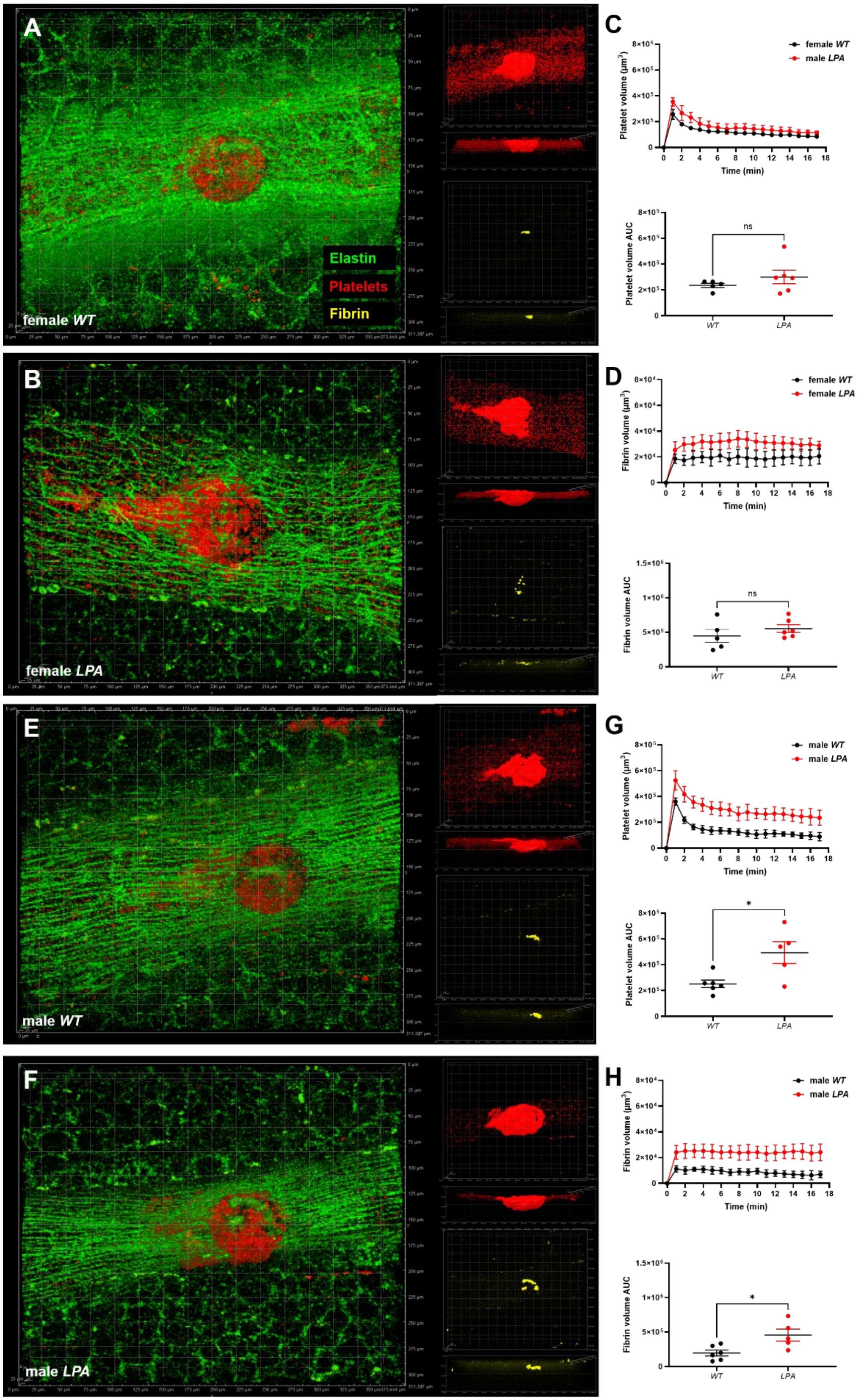
A
Elastin
Platelets
Fibrin
female WT
C
female WT
male LPA
Platelet volume (μm³)
Time (min)
ns
Platelet volume AUC
WT
LPA
B
female LPA
D
female WT
female LPA
Fibrin volume (μm³)
Time (min)
ns
Fibrin volume AUC
WT
LPA
E
male WT
G
male WT
male LPA
Platelet volume (μm³)
Time (min)
*
Platelet volume AUC
WT
LPA
F
male LPA
H
male WT
male LPA
Fibrin volume (μm³)
Time (min)
*
Fibrin volume AUC
WT
LPA

**Supplementary Fig. S4. Apo(a) increases platelet and fibrin volumes in standard diet-fed male mice in a laser-induced mesenteric vessel thrombosis model assessed by intravital imaging.** Thrombus formation in mesenteric veins was followed for a period of 17 minutes to determine platelet and fibrin volumes within developing thrombi. Representative XY and XZ 3D surface reconstructions of thrombi formed at t=1 min in (**A**) wild-type (*WT*); n=5) and (**B**) transgenic apo(a)-expressing (*LPA*; n=6) female mice fed standard diets. Overlay images contain colour composite green (emission wavelength: 488 nm; auto-fluorescent elastin), yellow (emission wavelength: 546 nm; fibrin), and red (emission wavelength: 647 nm; platelets). (**C**) Quantitative data displaying platelet volumes (upper panel) and scatterplots of distributions of cumulative platelet volume AUC (lower panel) in standard diet-fed female mice. (**D**) Quantitative data showing fibrin volumes (upper panel) and scatterplots of distributions of cumulative fibrin volume AUC (lower panel) in standard diet-fed female mice. Representative XY and XZ 3D surface reconstructions of thrombi formed at t=1 min are also shown for (**E**) male *WT* (n=6) and (**F**) male *LPA* (n=5) mice fed standard diets. (**G**) Quantitative data displaying platelet volumes (upper panel) and cumulative platelet volume AUC (lower panel) in standard diet-fed male mice. (**H**) Quantitative data showing fibrin volumes (upper panel) and cumulative fibrin volume AUC (lower panel) in standard diet-fed male mice. Quantitative data are shown as means ± SEM. Overlaid grid squares have dimensions of 25 × 25 $\mu m^2$. *$p < 0.05$ by Student’s t-test.

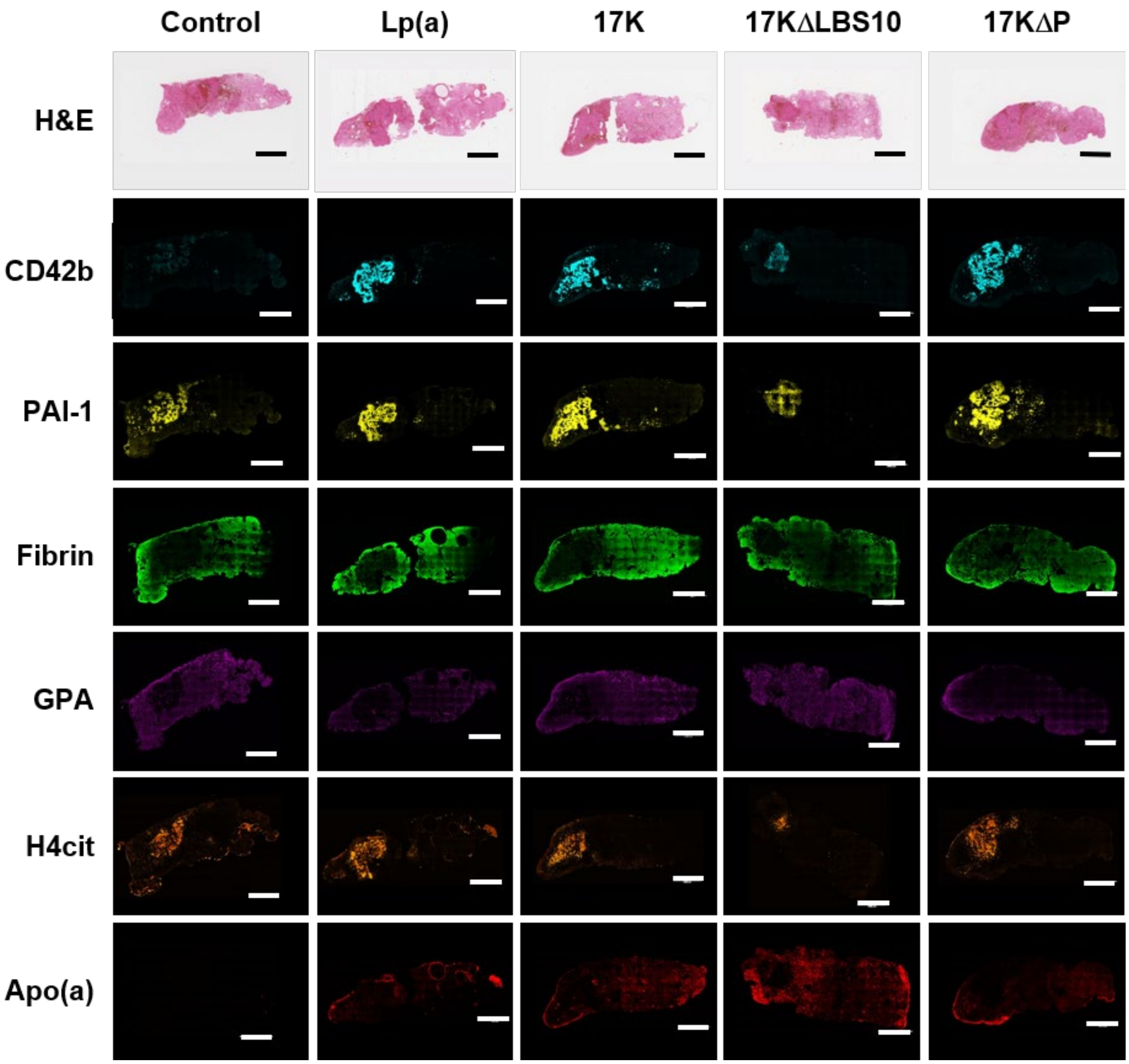


**Supplementary Fig. S5. Representative large-scan images of Chandler loop clots.** Clots were formed in the presence of HBS (control) or 250 nmol/L Lp(a), 17K apo(a), 17KΔLBS10, or 17KΔP. Following thrombogenesis in the Chandler loop apparatus, clots were immediately fixed in 4% paraformaldehyde, paraffin embedded, and subsequently sectioned longitudinally in 5-μm steps prior to Hematoxylin & eosin (H&E) and immunofluorescent (IF) staining. Clots were IF stained to detect the distribution of platelets (CD42b; cyan), plasminogen activator-1 (PAI-1; yellow), fibrin (green), red blood cells (GPA; magenta), neutrophil extracellular traps (H4Cit; orange) and apolipoprotein(a) (apo(a); red). Composite clot section images were acquired using the Nikon AIR+ confocal laser scanning system equipped with a 20× water-immersion objective. The 'Scan Large Image' tool was utilized to capture a representative image of the complete thrombus, and the XYZ overview panel allowed the software to interpolate and predict the appropriate focal height of all areas imaged on the slide. Colour composite and pseudocoloured images show emission at either 488 nm (CD42b, fibrin, GPA, H4Cit) or 647 nm (PAI-1, apo(a)). Images are shown at the same brightness/intensity scale for each ~~particular~~ antigen of interest. H&E sections were scanned using a Leica Aperio AT2 bright-field digital slide scanner with a 40× objective. Scale bars = 1000 μm.

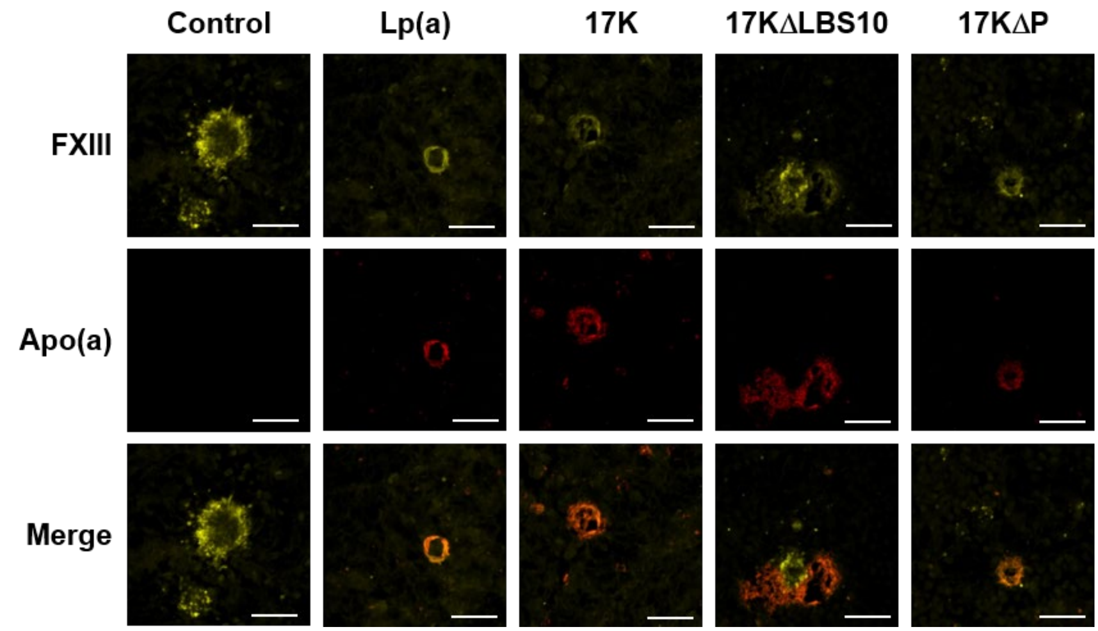


**Supplementary Fig. S6. Lp(a) and apo(a) appear to associate with fibrin pores through FXIIIa-mediated cross-linking.** Representative images of Chandler loop clot sections co-stained to detect the presence of apo(a) (red) and activated coagulation factor XIII (FXIIIa; yellow). Clots were formed in the presence of HBS (control) or 250 nmol/L Lp(a), 17K apo(a), 17KΔLBS10, or 17KΔP. Following thrombogenesis in the Chandler loop apparatus, clots were immediately fixed in 4% paraformaldehyde, paraffin embedded, and then ~~subsequently~~ sectioned longitudinally in 5-µm steps prior to immunofluorescent staining. Z-stack images of clot sections were acquired under identical acquisition settings and excitation wavelengths using the Nikon AIR+ confocal laser scanning system equipped with a 40× oil-immersion objective and high-speed Resonant scanner. Representative micrographs show z projections of z-stack image files (13 total z-slices, step size of 0.925 µm, 3× digital zoom). Images are shown at the same brightness/intensity scale. Overlay images are pseudocoloured yellow (emission at 647 nm) and red (emission at 488 nm).

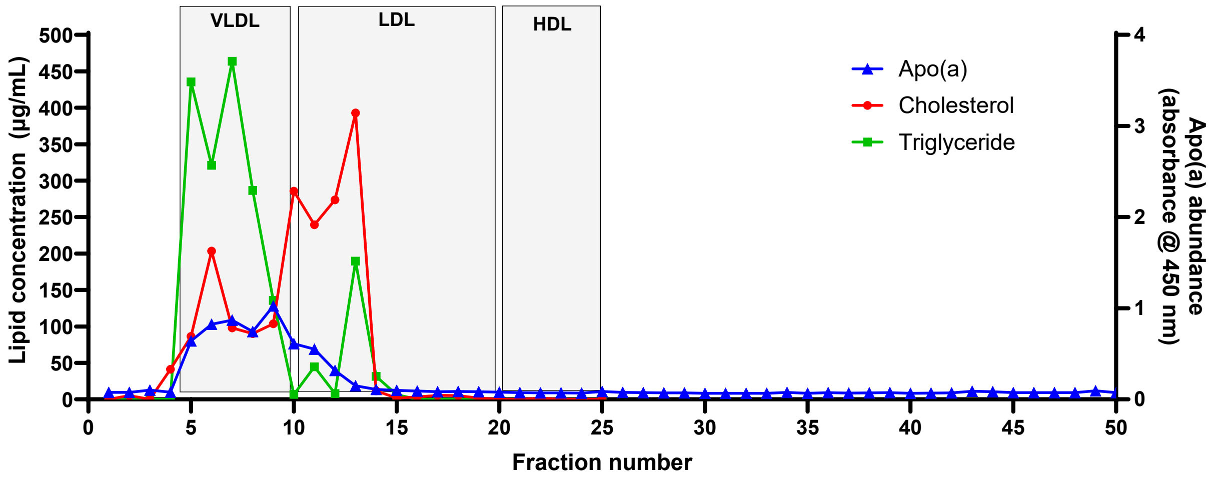


**Supplementary Fig. S7. Fast Protein Liquid Chromatography of plasma from *LPA* mice fed a HFHC diet.** Fresh plasma was passed over a Superose 6 size-exclusion column, and 500 µL fractions were collected for measurements of total cholesterol, triglycerides and apo(a). A total of fifty fractions were collected to detect the presence of any free apo(a) in plasma in later elution volumes. The expected elution fractions for different lipoprotein classes are indicated.

**Supplementary Table S1. Baseline characteristics of participants included in the UK Biobank analysis**

| Characteristic | **Without arterial thrombosis or VTE**, N = 433,490[1] | **With arterial thrombosis but no VTE**, N = 1,345[1] | **With VTE but no arterial thrombosis**, N = 10,644[1] |
|---|---|---|---|
| **Age** | 58 (50, 63) | 62 (57, 66) | 62 (56, 66) |
| **Male** | 197,056 (45%) | 924 (69%) | 5,568 (52%) |
| **Body Mass Index, Kg/m²** | 26.7 (24.1, 29.8) | 27.8 (24.8, 31.0) | 28.2 (25.5, 31.8) |
| **Waist circumference, cm** | 90 (80, 99) | 96 (87, 105) | 95 (86, 104) |
| **Diastolic blood pressure, mm Hg** | 82 (75, 89) | 82 (74, 89) | 83 (76, 90) |
| **Hypertension** | 171,830 (40%) | 1,032 (77%) | 6,030 (57%) |
| **Smoking** | | | |
| **Currently** | 44,846 (10%) | 426 (32%) | 1,302 (12%) |
| **Never** | 237,719 (55%) | 341 (25%) | 5,155 (48%) |
| **Prefer not to answer** | 1,714 (0.4%) | 9 (0.7%) | 54 (0.5%) |
| **Previously** | 149,211 (34%) | 569 (42%) | 4,133 (39%) |
| **Diabetes** | 21,769 (5.0%) | 246 (18%) | 824 (7.8%) |
| **Total Cholesterol, mmol/L** | 5.65 (4.91, 6.43) | 5.26 (4.43, 6.15) | 5.60 (4.82, 6.40) |
| **LDL-Cholesterol, mmol/L** | 3.52 (2.95, 4.12) | 3.25 (2.65, 3.94) | 3.50 (2.90, 4.12) |
| **HDL-Cholesterol, mmol/L** | 1.40 (1.17, 1.68) | 1.23 (1.03, 1.49) | 1.34 (1.13, 1.61) |
| **Triglycerides, mmol/L** | 1.48 (1.04, 2.14) | 1.70 (1.22, 2.52) | 1.62 (1.14, 2.29) |
| **Glycated hemoglobin, mmol/mol** | 35.2 (32.7, 37.8) | 37.7 (34.7, 41.9) | 36.1 (33.6, 39.0) |
| **Creatinine, umol/L** | 70 (61, 81) | 75 (65, 87) | 73 (64, 84) |
| **apoA-I, mmol/L** | 1.51 (1.35, 1.70) | 1.40 (1.25, 1.60) | 1.49 (1.32, 1.67) |

| | | | |
|---|---|---|---|
| **apoB, mmol/L** | 1.02 (0.86, 1.18) | 0.98 (0.81, 1.18) | 1.02 (0.86, 1.18) |
| **European ancestry, %** | 383,036 (88%) | 1,196 (89%) | 9,688 (91%) |
| **Lp(a), nmol/L** | 20 (8, 75) | 24 (8, 113) | 20 (8, 76) |
| **athromb** | 0 (0%) | 1,345 (100%) | 0 (0%) |
| **vte** | 0 (0%) | 0 (0%) | 10,644 (100%) |
| **Systolic blood pressure, mm Hg** | 136 (125, 149) | 143 (129, 157) | 139 (127, 151) |
| **Statins** | 76,145 (18%) | 613 (46%) | 2,665 (25%) |
| **Anti Hypertension Medication** | 88,255 (20%) | 647 (48%) | 3,127 (29%) |
| **Insulin** | 4,990 (1.2%) | 98 (7.3%) | 182 (1.7%) |
| **Coronary Artery Disease** | 13,129 (3.0%) | 230 (17%) | 491 (4.6%) |
| **Myocardial Infarction** | 9,505 (2.2%) | 156 (12%) | 364 (3.4%) |
| **Stroke** | 5,547 (1.3%) | 76 (5.7%) | 261 (2.5%) |

**Supplementary Table S2: Impact of lipoprotein(a) on the incidence of arterial embolism or thrombosis and venous thromboembolism in subsets defined by sex and age in the UK Biobank.** Cox proportional hazards for the study outcomes are shown for all participants in the UK Biobank, and for only men, only women, only participants less than 58 years old (median age) at baseline, and only participants greater than or equal to 58 years old (median age) at baseline.

| **Outcome** | **All (n=445,479)** | | **Men (n=203,548)** | | **Women (n=241,931)** | | **<58 years old (n=218,659)** | | **≥58 years old (n=226,820)** | |
|---|---|---|---|---|---|---|---|---|---|---|
| | HR (95% CI) | P | HR (95% CI) | P | HR (95% CI) | P | HR (95% CI) | P | HR (95% CI) | P) |
| **Arterial embolism or thrombosis** | 1.39 | 7.24E-07 | 1.50 | 3.13E-07 | 1.17 | 1.74E-01 | 1.34 | 2.64E-02 | 1.41 | 6.63E-06 |
| **Venous thrombo-embolism** | 1.03 | 2.60E-01 | 1.03 | 3.98E-01 | 1.03 | 4.80E-01 | 1.12 | 1.64E-02 | 1.00 | 8.83E-01 |
